# CG7379/ING1 suppress cancer cell invasion by maintaining cell-cell junction integrity

**DOI:** 10.1101/2020.08.20.255786

**Authors:** Alexandra D. Rusu, Zoe E. Cornhill, Brenda Canales Coutino, Marcos Castellanos Uribe, Anbarasu Lourdusamy, Zsuzsa Markus, Sean T. May, Ruman Rahman, Marios Georgiou

## Abstract

Approximately 90% of cancer related deaths can be attributed to a tumour’s ability to spread. We have identified CG7379, the fly orthologue of human ING1, as a potent invasion suppressor. ING1 is a type II tumour suppressor with well-established roles in the transcriptional regulation of genes that control cell proliferation, response to DNA damage, oncogene-induced senescence and apoptosis. Recent work suggests a possible role for ING1 in cancer cell invasion and metastasis, but the molecular mechanism underlying this observation is lacking. Our results show that reduced expression of CG7379 promotes invasion in vivo in *Drosophila*, reduces the junctional localisation of several adherens and septate junction components, and severely disrupts cell-cell junction architecture. Similarly, ING1 knockdown significantly enhances invasion in vitro and disrupts E-cadherin distribution at cell-cell junctions. A transcriptome analysis reveals that loss of ING1 affects the expression of several junctional and cytoskeletal modulators, confirming ING1 as an invasion suppressor and a key regulator of cell-cell junction integrity.

## Introduction

Metastasis is the major cause of mortality in human cancers. Underlying this phenomenon are a number of highly complex cellular behaviours, whereby cancer cells must acquire an ability to invade out of the primary tumour mass, avoid anoikis, migrate directionally, and disseminate to form secondary tumours at distant secondary sites. Unfortunately, the molecular mechanisms underlying these processes are poorly understood.

We have developed an in vivo system in *Drosophila* that allows the study of epithelial cell and tissue morphogenesis in real time (Georgiou et al., 2008, Georgiou and Baum, 2010, Cohen et al., 2010, Couto et al., 2017). We recently used this system to generate tumours with specific genotypes on the dorsal thorax epithelium of the fly and to screen for conserved modulators of tumour behaviour in the living animal (Canales Coutino et al., 2020). This screen allowed the identification of numerous invasion suppressors, one of which was the *Drosophila* inhibitor of growth (ING) orthologue CG7379.

CG7379 remains largely uncharacterised in *Drosophila*. Due to the presence of conserved zinc finger and plant homeodomain (PHD)-type domains (He et al., 2005) CG7379 has a putative role in chromatin remodelling and the transcriptional regulation of target genes. The human orthologue of CG7379 is ING1 and other ING family members (Flockhart et al., 2012). In mammals there are five members of theING family of proteins (ING1-5, including several splice variants), and virtually all members of this family have been shown to possess tumour suppressive functions (Jafarnejad and Li, 2012). All ING family members contain the signature C-terminal PHD finger domain. This highly conserved domain has the highest degree of homology among the ING proteins (He et al., 2005) and also shares a very strong identity of 78% with the PHD domain of CG7379 (Flockhart et al., 2012).

Loss of chromosomal locus, translocation to the mitochondria, mutation (rarely), but mainly decreased expression of ING1 have been documented in various mammalian cancers (Ythier et al., 2008, Bose et al., 2014). Several studies investigating the effect of ING1 on cell proliferation, reported cells accumulating in G0/G1 upon ING1b overexpression in a range of cell lines from normal fibroblasts to cancer cells derived from metastatic sites (Abad et al., 2011, He et al., 2014, Jia et al., 2015, Jiang et al., 2018). ING1 is a key player in multiple DNA repair pathways (Cheung and Li, 2002, Garate et al., 2007, Ceruti et al., 2013), with evidence suggesting that ING1 can induce apoptosis in response to DNA damage (Thalappilly et al., 2011, Russell et al., 2006). The antiproliferative and proapoptotic effects of ING1 were linked to its ability to modulate transcription. ING1 physically interacts with protein complexes with histone acetyltransferase (HAT) and histone deacetylase (HDAC) activity (Russell et al., 2006). ING1b is a stable component of the Sin3A/HDAC 1 and 2 protein complexes and can additionally interact with subunits of the Brg1-based Swi/Snf chromatin remodelling complex (Kuzmichev et al., 2002a, Peña et al., 2008, Kuzmichev et al., 2002b), with Proliferating Cell Nuclear Antigen (PCNA) and p300 (Russell et al., 2006). Additionally, ING1 has been shown to modulate expression of several micro RNAs, which in turn further regulate gene expression (Chen et al., 2013, He et al., 2014).

Several studies have also associated ING family proteins with invasion and metastasis. For example, reduced expression of ING1b and ING4 has been observed in metastatic melanoma (Nouman et al., 2002, Li et al., 2008) and reduced ING4 expression could be correlated with human gastric adenocarcinoma stage (Li et al., 2009). Low levels of ING1 are associated with increased motility, migration or invasion in colorectal, gastric and breast cancers (Ahmed et al., 2001, Guo et al., 2011, Thakur et al., 2014). It was further shown that ING4 overexpression inhibited melanoma cell migration and invasion in vitro (Li et al., 2008). ING1 overexpression was also shown to inhibit cell migration, invasion and metastasis both *in vitro* and *in vivo* (Thakur et al., 2014, Jiang et al., 2018) and ING1 knock down in MDA-MB-231 cells increased migration and invasion (Thakur et al., 2014). However, despite ING family proteins being repeatedly implicated in tumour progression and invasion, little is known of the molecular mechanisms that underlie these tumour suppressive effects. Here, using the *Drosophila* ING orthologue CG7379, we demonstrate that loss of this protein results in a severe disruption to the adherens and septate junctions, resulting in a loss of epithelial integrity and increased invasion. We further show that loss of ING1 in human cancer cells also promotes invasion and disrupts cell-cell adhesion, through altered gene expression.

## Results

### CG7379 acts as an invasion suppressor

We recently carried out an in vivo large-scale screen for genes that affect tumour behaviour by: (1) generating positively marked clones on the dorsal thorax of the fly that are homozygous mutant for the neoplastic tumour suppressor gene *lethal (2) giant larvae (lgl*^*4*^); (2) specifically labelling the mutant tissue with GFP:Moe (the actin binding domain of moesin fused to GFP), thereby labelling the actin cytoskeleton of these cells; (3) overexpressing an RNAi transgene to deplete expression of a gene of interest specifically within the mutant, labelled tissue. In this way, we could identify genes that when knocked down (KD) work cooperatively with *lgl*^*4*^ to promote tumour progression (Canales Coutino et al., 2020).

Our screen identified CG7379 as a strong hit for invasion, with *lgl*^*4*^; CG7379KD mutant clones showing a significant increase in the number of polarised epithelial cells seen beneath the epithelial sheet ((Canales Coutino et al., 2020); Figure 1a-e). As well as promoting frequent cell delamination, we found *lgl*^*4*^; CG7379KD clones to be multilayered and to promote abnormal protrusion morphology (Figure S1).

**Figure 1:**
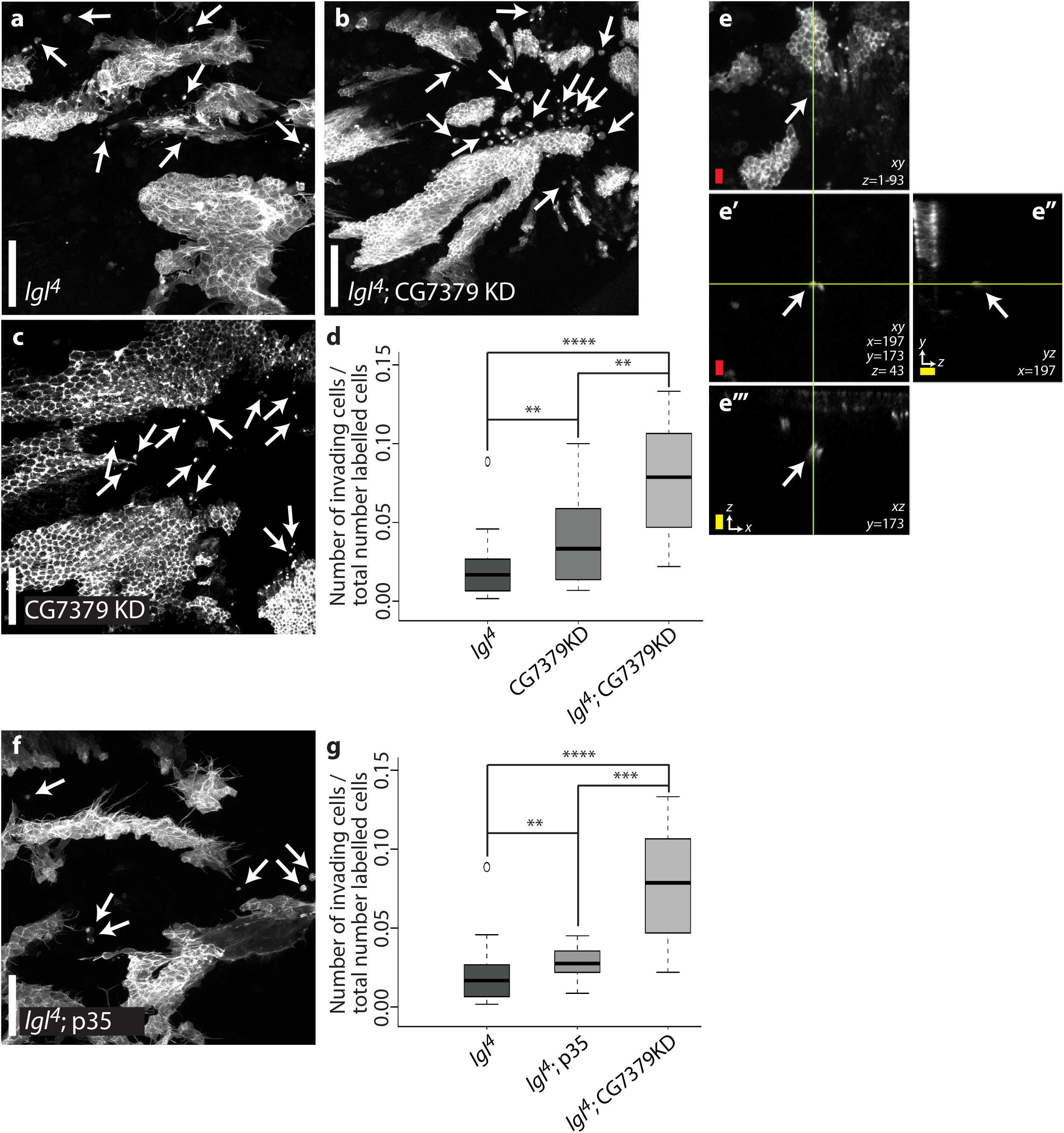
CG7379 KD promotes invasion. **(a-d)** GFP:Moe labelled clones mutant for (a) *lgl*^*4*^, (b) *lgl*^*4*^; CG7379KD, and (c) CG7379KD. CG7379KD results in a highly invasive phenotype, which is enhanced when accompanied by a *lgl*^*4*^ mutation (quantified in d). **(e)** z max intensity projection of the back of a 28h APF (after puparium formation) old *Drosophila* pupa carrying GFP:Moe labelled CG7379KD clones. **(e’)** Individual confocal z slice highlighting a GFP:Moe labelled cell located 25µm beneath the epithelial sheet. z = confocal slice; x and y = coordinates and position of the invasive cell. **(e’’)** yz orthogonal view projection of the invasive cell in (e’). **(e’’’)** xz orthogonal view projection of the invasive cell in (e’). **(f-g)** *lgl*^*4*^; p35 (f) does not phenocopy *lgl*^*4*^; CG7379KD (b). Although a significant increase in invading cells is observed when overexpressing p35 within *lgl*^*4*^ clones, this increase is significantly lower than that observed in *lgl*^*4*^; CG7379KD clones (quantified in g). *lgl*^*4*^ n=42, CG7379KD n=21, *lgl*; CG7379KD n=21, *lgl*; p35 n=21). Kruskal-Wallis test was performed to determine statistical significance. White scale bar: 50µm, red scale bar: 10µm, yellow scale bar: 10µm in the z plane. *p<0.05; **p<0.01; ***p<0.001 ****p<0,0001.

GFP:Moe labelled CG7379KD clones, without the *lgl*^*4*^ mutation, formed an organised monolayered epithelium with significantly fewer invading cells found beneath the epithelium when compared to *lgl*^*4*^; CG7379KD mutant clones (Figure 1b-d). However, there was also a significant increase in invasion when compared to *lgl*^*4*^ mutant clones (Figure 1d). It therefore appears that although CG7379KD promotes epithelial cell delamination, there is a strong cooperative effect when this KD is accompanied by a loss of lgl function, leading to increased multilayering and invasion.

### Increased invasion following CG7379KD is in part due to cell death evasion

It is known that ING1 can act as a tumour suppressor by inducing apoptosis in damaged cells (Russell et al., 2006, Bose et al., 2013). It is therefore possible that the invasive phenotype that we observe when knocking down CG7379 in the *Drosophila* notum could be due to the increased survival of invading cells, whereby invading cells are able to evade elimination via apoptosis. To test this, we overexpressed P35 specifically within GFP:Moe labelled *lgl*^*4*^ mutant clones. Overexpression of P35 is known to prevent apoptotic death in a wide variety of tissues and is widely used in *Drosophila* cell death studies (Hay et al., 1994). If the increase in invasion observed in *lgl*^*4*^; CG7379KD mutant clones was largely due to an inhibition of apoptosis, we would expect to see a similar increase in GFP:Moe positive cells beneath the epithelial sheet when expressing P35 in *lgl*^*4*^ mutant clones. This however was not observed. Although a significant increase in GFP:Moe positive cells was observed when comparing *lgl*^*4*^ mutant clones with *lgl*^*4*^; P35 clones (P=0.009), we did not see the dramatic increase in the number of invasive cells that we observe in lgl^4^; CG7379KD clones (Figure 1f-g).

We additionally looked for apoptosis within pre-invasive cells, using both a TUNEL assay and an anti-cleaved *Drosophila* Dcp-1 antibody (Figure 2a-b). Pre-invasive cells are cells that are about to delaminate from the epithelial sheet and can be identified by their rounded morphology and the presence of a characteristic actin-rich spot at one side of the cell prior to invasion (Canales Coutino et al., 2020). CG7379KD simultaneously significantly increased the number of pre-invasive cells present within a clone, and also reduced the amount of apoptosis occurring within these pre-invasive cells, when compared to WT clones. However, when comparing *lgl*^*4*^; CG7379KD clones with *lgl*^*4*^ clones, no effect on the proportion of apoptotic pre-invasive cells was observed (Figure 2b). This suggests that although CG7379KD most likely plays a role in preventing apoptosis, the significant increase in the number of invading cells following CG7379KD is unlikely to be only due to cell death evasion.

**Figure 2:**
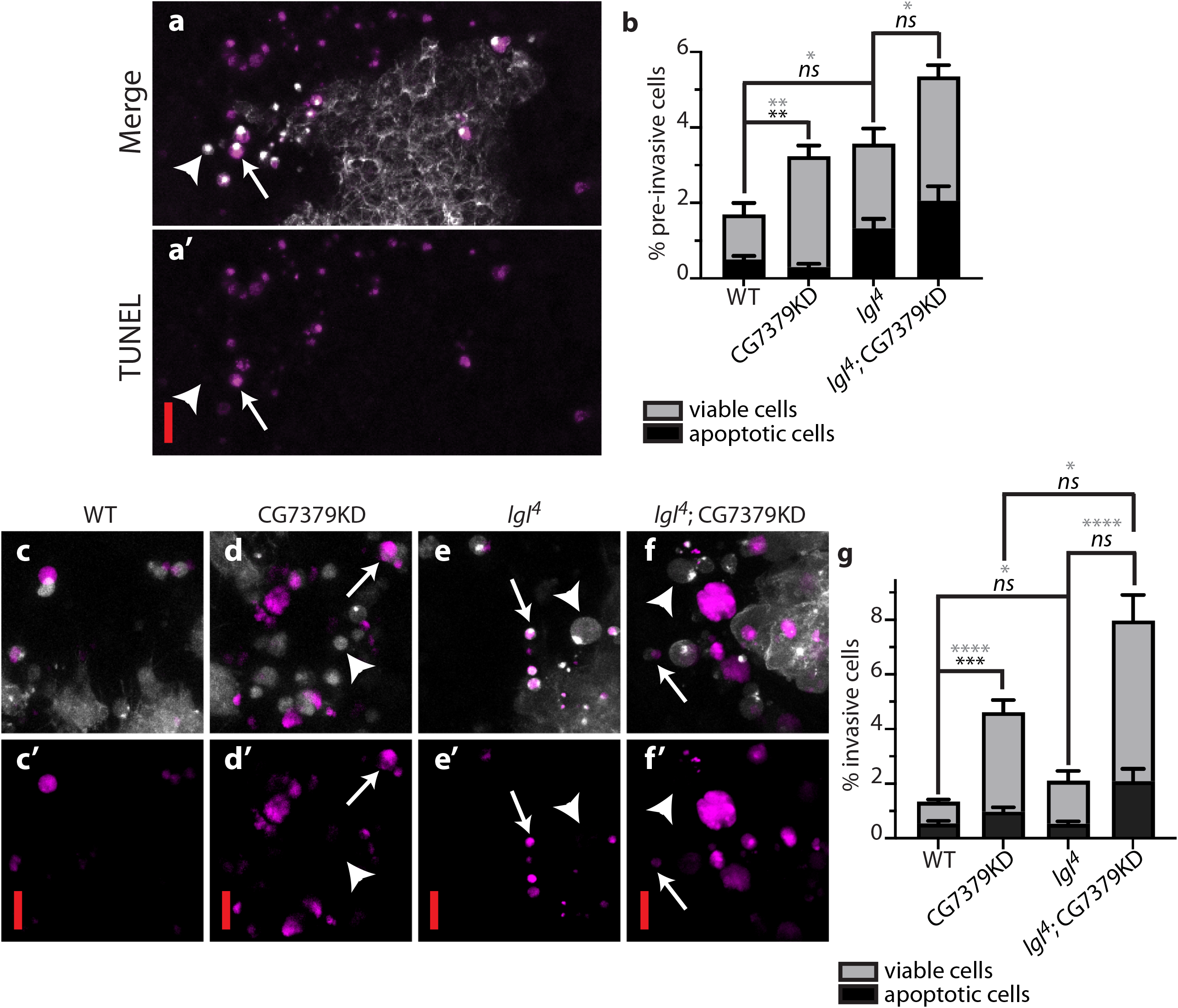
CG7379KD promotes evasion of apoptosis. **(a)** A TUNEL (magenta) and GFP:Moe (white) labelled *lgl*^*4*^ mutant clone, highlighting both apoptotic (arrow) and non-apoptotic (arrowhead) pre-invasive cells. **(b)** Quantification of the proportion of apoptotic pre-invasive cells (black), as determined by TUNEL and c-Dcp-1 staining. Grey, % pre-invading cells/total number of labelled cells; black, % apoptotic pre-invading cells/total number of labelled cells. When comparing CG7379KD with WT clones, significantly more pre-invasive cells (grey asterisks) were observed, out of which significantly fewer were apoptotic (black asterisks) (n≥138 cells across 10 animals/genotype). **(c-f)** Example images of apoptotic (arrow) and non-apoptotic (arrowhead) invasive cells. iCasper (magenta) and Moe:GFP (white) labelled mutant clones (genotypes specified above panels). Images represent z stack projections. **(g)** Quantification of (c-f). Grey, % invading cells/total number of labelled cells; black, % apoptotic invading cells/total number of labelled cells. n≥346 cells across 20 animals/genotype. Kruskall-Wallis test was performed to determine statistical significance. Error bars represent ± s.e.m. Scale bars: 10µm. *p<0.05; **p<0.01; ***p<0.001 ****p<0,0001.

The viability of cells once they had delaminated from the epithelial sheet was next assessed using the in vivo caspase sensor, iCasper (Figure 2c-g) (To et al., 2015). We found the proportion of apoptotic invasive cells was significantly reduced by CG7379KD when compared to WT clones. However, although *lgl*^*4*^; CG7379KD dramatically increases the number of viable invading cells when compared to *lgl*^*4*^ clones, no significant reduction in the proportion of apoptotic invasive cells was observed (Figure 2g). These results provide further evidence to suggest that the cooperative effect observed between the *lgl*^*4*^ and CG7379KD mutations is not simply due to an increase in cell viability.

### CG7379 is required to maintain epithelial cell-cell junction integrity

Reduced ING expression has been previously observed in high-grade metastatic cancers (Nouman et al., 2002, Li et al., 2008, Li et al., 2009). However, despite ING family proteins being implicated in invasion and metastasis, little is known of the molecular mechanisms that underlie these effects.

Our screen identified *lgl*^*4*^; CG7379KD as affecting epithelial architecture, protrusion morphology and promoting frequent cell delamination ((Canales Coutino et al., 2020); Figures 1 and S1). These phenotypes therefore implicate effects on adhesion, polarity and actin regulation as possible underlying influences on the observed cell behaviour. To investigate the molecular mechanisms by which loss of CG7379 expression could affect epithelial cell organisation, we first investigated cell-cell adhesion using antibodies to proteins that localise to either the adherens junction (AJ) or septate junction (SJ, the functional equivalent of tight junctions in insects). We stained for E-cadherin, Armadillo (β-catenin), and α-catenin (AJ proteins), and Fasciclin III, Coracle and Discs Large (SJ proteins) (Figures 3 and S2). We generated positively marked mutant clones, surrounded by wild-type tissue, thereby allowing us to directly compare junction composition inside and outside mutant clones in the same tissue. We looked in mutant clones for *lgl*^*4*^; CG7379KD, CG7379KD alone and *lgl*^*4*^ alone. In most cases, AJ and SJ protein localisation was significantly reduced, and AJ and SJ integrity was significantly and severely disrupted, in both *lgl*^*4*^; CG7379KD and CG7379KD clones, with minor effects observed in *lgl*^*4*^ clones (Figure 3). The only exception was when staining for Dlg, where the most severe effects were observed in *lgl*^*4*^ mutant clones (Figure S2a-d).

**Figure 3:**
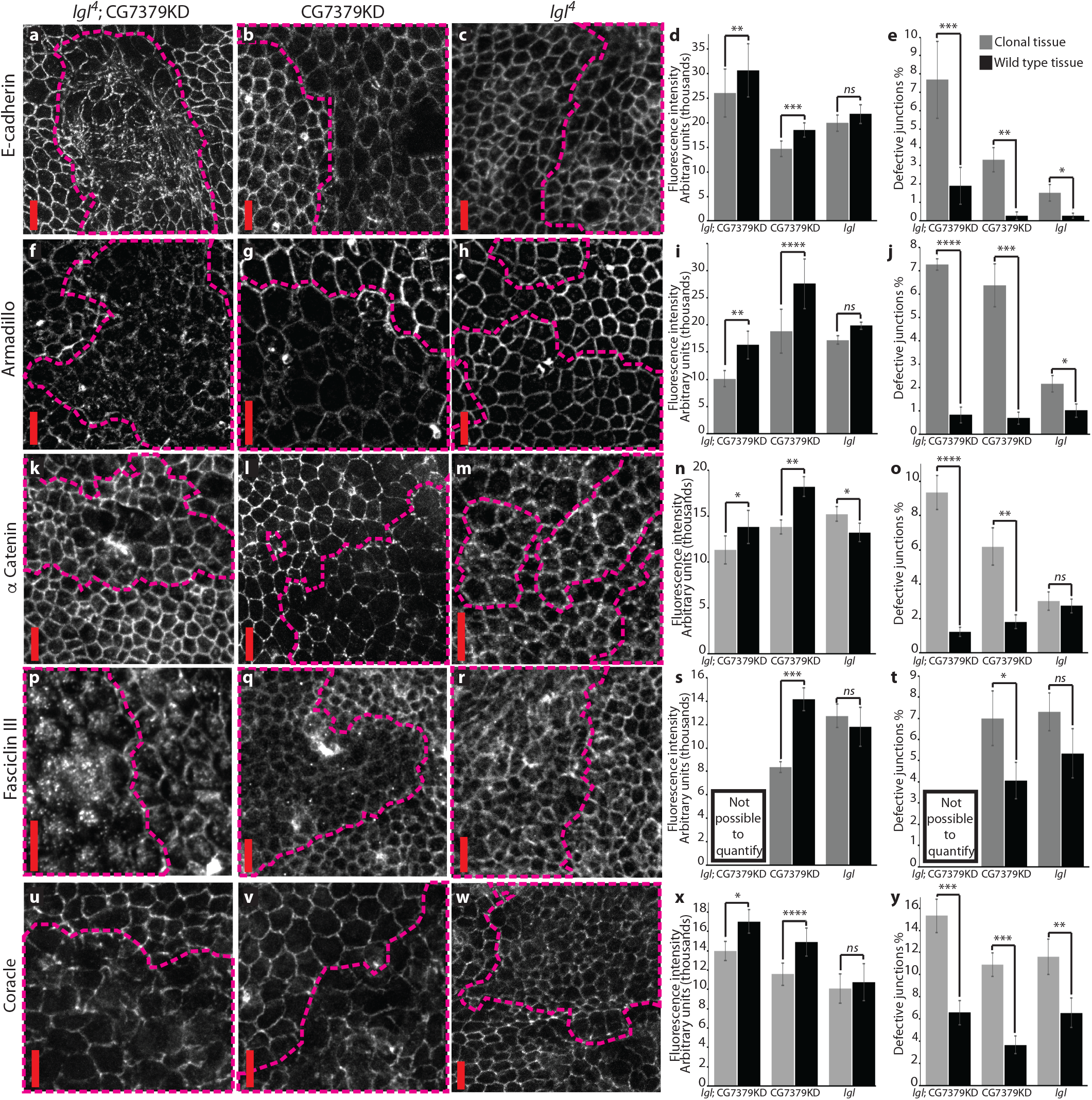
CG7379KD disrupts cell-cell junction integrity. *Drosophila* pupal nota containing positively marked mutant clones for *lgl*^*4*^; CG7379KD; CG7379KD alone; or *lgl*^*4*^ alone (highlighted by dashed lines). Nota were stained for the following junctional proteins: **(a-c)** E-cadherin, **(f-h)** Armadillo (β-catenin), **(k-m)** α-Catenin, **(p-r)** Fasciclin III, **(u-w)** coracle. A severe disruption in junction protein localisation and junction integrity was observed in *lgl*; CG7379KD and CG7379KD clones, as quantified in panels **(d-e; i-j; n-o; s-t; x-y)** (n≥ 65 junctions from a minimum of 9 animals for each genotype). Fasciclin III staining could not be quantified in *lgl*; CG7379KD clones due to an almost complete lack of protein localisation at the cell cortex. Scale bars = 10 μm. Error bars represent ± s.e.m. Student’s T test was performed to determine statistical significance. *p<0.05; **p<0.01; ***p<0.001 ****p<0,0001.

We next looked at whether the apicobasal polarity determinants aPKC, bazooka (Par3) or Crumbs were affected in these mutant clones, as a disruption of apicobasal polarity frequently promotes invasion. We only found a significant effect on aPKC localisation, which is likely due to a loss of lgl function, as no significant effect was found in CG7379KD clones (Figure S2). The additional effects on cell polarity in *lgl*^*4*^mutant clones (i.e. the mislocalisation of both Dlg and aPKC) likely explain the increased multilayering and invasion observed in *lgl*^*4*^; CG7379KD clones, when compared with CG7379KD alone.

Since E-cadherin is at the core of cell-cell junction adhesion and since E-cadherin localisation at the AJ was severely disrupted in both *lgl*^*4*^; CG7379KD and CG7379KD clones, we hypothesised that if E-cadherin levels could be restored, this may partially rescue cell-cell junction defects and consequently cell delamination. Therefore, we attempted to rescue the CG7379KD invasive phenotype by overexpressing E-cadherin under the Ubi-p63E promoter in CG7379KD mosaic animals (Figure S3). The construct we used has been shown to substitute for a null *shotgun* allele (the gene encoding E-cadherin in flies) and to function and behave normally in the absence of intact E-cadherin (Oda and Tsukita, 2001). We however found no significant effect on invasion, suggesting that either (1) E-cadherin overexpression is not sufficient to compensate for the CG7379KD-mediated disruption to multiple AJ and SJ junction components; or (2) E-cadherin mis-localisation to the AJ, rather than simple expression levels, may be responsible for the observed phenotype.

### ING1 regulates cell adhesion through transcriptional regulation of cell adhesion modulators

Having identified CG7379 as an important invasion suppressor, we wanted to test whether its closest human orthologue, ING1, would also act in a similar way. Using an antibody against human E-cadherin on the breast cancer cell line MCF7, we found a significant disruption to both E-Cadherin localisation and to AJ integrity following ING1KD, with frequent junctional breaks observed (Figure 4a-c), suggesting that human ING1 also plays an important role in maintaining AJ integrity. We recently demonstrated that KD of ING1 led to an increase in migration and invasion in MCF7 cells, using an in vitro invasion assay (Canales Coutino et al., 2020). Here, we additionally show that this effect on invasion is not restricted to MCF7 cells, as ING1KD also increases invasion in U87 (derived from malignant glioma) and MDA-MB-231 cells (derived from malignant breast adenocarcinoma) (Figure 4d-f). Relative ING1 mRNA levels following KD, as determined by qRT-PCR were 28% (MCF7), 4% (MDA-MB-231), and 9% (U87) (data not shown). In line with these results, ING1KD has previously been shown to increase migration and invasion in MDA-MB-231 cells (Thakur et al., 2014).

**Figure 4:**
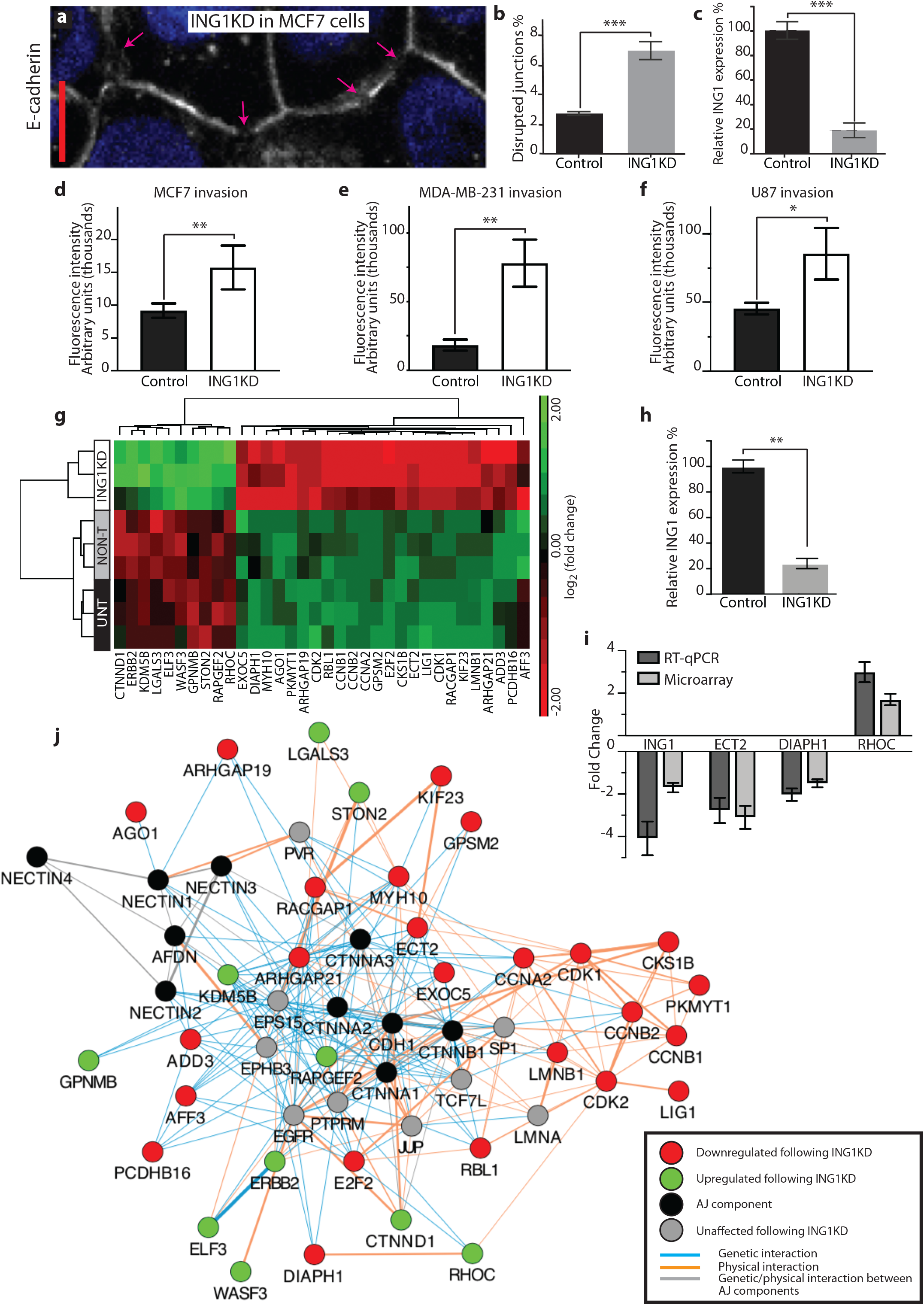
ING1KD increases the invasive potential of several cancer cell lines and disrupts cell-cell junction integrity in MCF7 cells. **(a-c)** MCF7 cells transfected with siRNA for ING1 and stained for E-cadherin. Disruption to E-cadherin localisation at the junction was observed (a, arrows highlight junctional breaks). Quantification shows a significant increase in disrupted junctions upon ING1KD (b). NON-T MCF7 cells were used as a control. n = 606 junctions for ING1KD and 823 junctions for the control. Relative ING1 mRNA levels, determined by qRT-PCR (c). **(d-f)** Transwell invasion assays for MCF7 (d), MDA-MB-231 (e), U87 cells (f). In order to invade, cells had to pass through an 8µm pore membrane coated with Matrigel. A significant increase in invasion for all three cell lines was observed following ING1KD in comparison to NON-T cells. **(g)** Heat map representation of unsupervised clustering of 34 selected differentially expressed genes following ING1KD in MCF7 cells. Columns represent genes, each row represents a sample. Treatment groups are specified on the left-hand side. Colour code represents log2 of the fold change of expression: red, downregulated; green, upregulated. Horizontal and vertical clusters were created based on Euclidean distance. **(h)** Relative ING1 mRNA levels, determined by qRT-PCR, for transcriptomics experiment. **(i)** Bar chart comparing microarray and RT-qPCR for four genes with differential expression between ING1KD cells and NON-T cells. **(j)** Interaction network showing physical and genetic interactions between selected differentially expressed genes by ING1KD and proteins associated with the AJ. The GeneMANIA plug-in for Cytoscape was used to generate an interaction network based on previously documented interactions. Black nodes mark AJ components, red nodes: genes that were downregulated by ING1KD; green nodes: genes that were upregulated by ING1KD; grey nodes: genes through which they interact (standard settings: max resultant attributes=10; max resultant genes=20); orange lines: physical interactions; blue lines: genetic interactions; grey lines genetic or physical interactions between AJ components. Line thickness is indicative of the score (weighting: GO molecular function). Scale bar = 10 μm. Error bars represent ± s.e.m. Student’s T test was performed to determine statistical significance. *p<0.05; **p<0.01; ***p<0.001 ****p<0,0001.

ING1 is known to act as both a transcriptional activator and repressor through interactions with DNA and a variety of epigenetic regulatory proteins (Coles and Jones, 2009). In an effort to understand how loss of ING1 function might lead to a disruption to cell-cell junctions we performed a microarray gene expression analysis. We analysed and compared gene expression in MCF7 cells post ING1KD with untreated cells (UNT) and cells treated with non-targeting siRNA (NON-T). Out of 21448 genes analysed, the expression of 919 genes was significantly altered as a result of ING1KD (FDR<0.05, p<0.01, FC≥1.5 or FC≤-1.5; Table S1; Figure S4). GO-enrichment analysis and pathway analysis showed that most of the significantly altered genes are involved in cell cycle regulation and DNA replication (Table S2) which corroborates previous work on ING1 function (Ythier et al., 2008). However, we additionally found 34 differentially expressed genes (DEGs) that are likely to affect both cell-cell junction integrity and promote invasive phenotypes, including DIAPH1, ECT2, EXOC5, WASF3, RAPGEF2, ADD3, KDM5B, CTNND1, and RHOC (Figure 4g). There is extensive evidence linking the misregulation of these genes with a disruption to cell-cell junction integrity, and to the promotion of invasion (see Discussion). We additionally used RT-qPCR on a selection of genes (RHOC, DIAPH1, ECT2, ING1) to verify the microarray results, with qPCR showing a similar or greater expression fold change in all cases (Figure 4i); results from the two analyses show high correlation (Pearson test: r=0.89, p=0.05).

The Cytoscape plugin GeneMANIA (Warde-Farley et al., 2010) was used to assess interactions between the selected subset of 34 DEGs and AJ components. For each selected DEG, we searched for physical or genetic interactions, validated by experimental data, including yeast two-hybrid, co-immunoprecipitation, and other interaction data from various databases (see Methods). The resultant network (Figure 4j; Data S1) reveals a total of 141 genetic interactions and 116 physical interactions between 34 DEGs and 10 AJ KEGG pathway genes (Table S3) indicating that the differentially expressed genes in ING1KD cells extensively interact with members of the AJ pathway.

## Discussion

The in vivo model for epithelial cancer that we have developed has proven to be particularly suitable for the study of cancer cell invasion. Major advantages of our system include our ability to generate tumours surrounded by wild-type tissue and the native local microenvironment, thereby maintaining the complex tumour-stroma interactions known to influence tumour behaviour (Valkenburg et al., 2018), and the ability to image tumour behaviour in real time in the living animal. This has led to the identification of numerous invasion suppressors (Canales Coutino et al., 2020), including CG7379, whose KD very strongly promotes invasion from the dorsal thorax epithelium. Although CG7379 is largely uncharacterised in *Drosophila*, it has been picked up in numerous screens, focusing on diverse biological functions, including Notch pathway regulation, muscle morphogenesis, adiposity regulation, and airway morphogenesis (Saj et al., 2010, Mummery-Widmer et al., 2009, Schnorrer et al., 2010, Baumbach et al., 2014, Hosono et al., 2015). The latter screen is important as it implicates CG7379 in the maintenance of epithelial integrity in the tubes that make up the fly tracheal system.

The human orthologues of CG7379, the ING family proteins, have been extensively implicated in invasion and metastasis but little was known of the molecular mechanisms that underlie these effects. Here, using the *Drosophila* ING orthologue CG7379, we demonstrate that loss of this protein results in a severe disruption to both the AJ and SJ, resulting in a loss of epithelial integrity and increased invasion. Results further suggest that loss of CG7379 also promotes cell survival following cell detachment from the epithelial sheet, demonstrating the multiple tumour and invasion suppressive roles of this gene.

Our transcriptomics experiments, knocking down ING1 expression in MCF7 breast cancer cells, provides some mechanistic insight into how loss of ING1 function might affect cell-cell junctions and promote invasion. The expression of numerous genes that regulate cell-cell adhesion and/or the actin cytoskeleton were altered following ING1KD in MCF7 cells. This included EXOC5, which was significantly downregulated and shows high homology to *Drosophila* sec10. Sec10 has been shown to interact with Armadillo (β-catenin) and α-Catenin (Langevin et al., 2005) and via sec5 and 15 can reduce E-cadherin, Armadillo and α-Catenin levels at AJs in *Drosophila* (Langevin et al., 2005). Furthermore, increased EXOC5 mRNA levels were previously correlated with E-cadherin overexpression within the lymphovascular embolus of inflammatory breast cancer (Ye et al., 2010).

RHOC and KDM5B were both significantly upregulated in our study. RHOC has been shown to be a marker of metastatic potential in some breast cancers (Wu et al., 2010). RHOC overexpression has been associated with increased cell motility and invasion through stress fibre formation and focal adhesion, whilst its KD has been shown to decrease invasion and nuclear β-catenin levels (Ridley, 2013, Simpson et al., 2004, Wu et al., 2010). An increase in RHOC expression was reported to induce a decrease in E-cadherin levels at the AJ in MCF7 cells (but not overall E-cadherin levels) without affecting Snail nor Twist levels nor promoting nuclear β-Catenin accumulation (Kawata et al., 2014). These results fit both our transcriptomics results, and the effect we observe on junction integrity in *Drosophila* nota when knocking down CG7379 expression.

In breast cancer, KDM5B and E-cadherin present reverse patterns of expression (Bamodu et al., 2016). Moreover, ectopic expression of KDM5B has been shown to promote EMT by downregulating E-cadherin through SNAIL independent mechanisms (Bamodu et al., 2016). Overexpression of KDM5B can also increase invasion in vitro and metastatic potential of gastric tumours in vivo, by activating the Akt pathway. KDM5B is therefore very likely to modulate cytoskeleton dynamics (Wang et al., 2014). In *Drosophila* haemocytes, little imaginal discs (lid), the fly orthologue of KDM5B, modulates Rac and Ras expression and consequently actin cytoskeleton organisation. Lid depletion has been shown to trigger lamellipodia formation and hyper-polymerisation of F-actin (Morán et al., 2015).

Adducin 3 (ADD3) expression was two-fold decreased in our study. Adducin proteins are critical for proper formation and stabilisation of the membrane cytoskeleton and regulation of cell motility and cell-cell adhesion (Luo and Shen, 2017). ADD3 depletion was shown to negatively affect both AJ and tight junction reassembly and to impair the assembly of actin filaments associated with newly formed junctions in colon epithelial cells (Naydenov and Ivanov, 2010). Similarly, in *Drosophila* embryos, hu-li tai shao (hts), the fly orthologue of human adducins, was shown to partially colocalise with dlg and to regulate dlg targeting to the membrane. Moreover, hts mutants displayed phenotypes indicative of disruptions to epithelial integrity (decreased cuticle secretion in embryos) (Wang et al., 2011). In *Drosophila* oocytes, improper localisation of hts RNA led to the overgrowth of actin filaments (Pokrywka et al., 2014).

DIAPH1, a downstream target of RHOA, was downregulated by ING1KD. Its loss has been associated with a decrease in the localisation of E-cadherin, α- and β-catenin at AJs, and also affected tight junction composition and junctional actin levels (Carramusa et al., 2007). Additionally, ING1KD also affected the expression levels of several regulators of Rho GTPase activity, including ECT2, a RhoGEF that is required to activate Rho signalling at the zonula adherens and support junctional integrity through myosin IIA (Ratheesh et al., 2012).

Our transcriptomics results therefore confirm ING1 as an invasion suppressor that plays an important role in maintaining junction stability, and as a modulator of cytoskeletal dynamics. This perfectly recapitulates results from the fly, where CG7379KD strongly promotes invasion, severely disrupts cell-cell junction integrity and affects protrusion morphology. The ING family of tumour suppressors are well known to suppress tumourigenesis through the regulation of cell cycle progression, apoptosis, and DNA repair. This work, by recognising CG7379 and ING1 as invasion suppressors, adds further insight into the multiple roles that collectively perform ING’s tumour suppressive function.

## Acknowledgments

We wish to thank the fly community for their generosity with reagents, especially the Wodarz lab for Baz antibody and the Bloomington, VDRC and NIG stock centres. We thank Anna Grabowska for the MCF7 cell line and School of Life Sciences Imaging (SLIM) for invaluable help with the confocal microscopes. This work was supported by Cancer Research UK [grant numbers C36430, A12891]. B.C.C. was supported by a CONACyT award; A.D.R. was supported by a Nottingham Vice-Chancellor’s Scholarship for Research Excellence Award.

## Author Contributions

Conceptualisation, M.G.; Methodology, M.G., R.R., A.D.R., B.C.C., and Z.E.C.; Investigation, A.D.R, B.C.C, Z.E.C., and Z.M.; Formal Analysis, A.L., M.C.U., A.D.R. and B.C.C.; Writing – Original Draft, M.G. and A.D.R; Writing – Review & Editing, M.G., A.D.R., R.R. and B.C.C. Funding Acquisition, M.G.; Resources, M.G., R.R., and S.T.M.; Supervision, M.G., R.R., and S.T.M.

---

**Figure S1: Abnormal protrusion morphology following CG7379KD**

**(a-c)** Example of typical basal protrusion morphology from *lgl*^*4*^ cells (a), and examples of abnormal branched protrusions from *lgl*^*4*^; CG7379KD (b) and CG7379KD (c) cells. Images represent z stack projections of relevant basal confocal slices. Arrows highlight branched protrusions. **(d)** Quantification of branched protrusion occurrence. n≥100 protrusions/ animal from a minimum of 7 animals/genotype. One-way ANOVA was performed to determine statistical significance. Scale bar = 10 μm. * p<0.05; **p<0.01; ***p<0.001; ****p<0.0001.

**Figure S2: CG7379KD does not affect the localisation of apicobasal polarity determinants**

*Drosophila* pupal nota containing positively marked mutant clones for *lgl*^*4*^; CG7379KD; CG7379KD alone; or *lgl*^*4*^ alone (highlighted by dashed lines). Nota were stained for the following proteins: **(a-c)** discs large, **(e-g)** aPKC, **(i-k)** bazooka, **(m-o)** crumbs. Discs large and aPKC disruption was most severe in *lgl*^*4*^ mutant clones, as quantified in panels **(d and h)**. No effect was observed when staining for the apical polarity proteins bazooka and crumbs **(l and p)**. n≥ 55 junctions from a minimum of 4 animals for each genotype. Scale bars = 10 μm. Error bars represent ± s.e.m. Student’s T test was performed to determine statistical significance. *p<0.05; **p<0.01; ***p<0.001 ****p<0,0001.

**Figure S3: E-cadherin overexpression cannot rescue CG7379KD induced cell delamination**

**(a-b)** Representative confocal image of GFP:Moe labelled CG7379KD clones in an otherwise WT *Drosophila* notum (a) and CG7379KD in a notum overexpressing E-cadherin (b). Images represent max projection of confocal z stacks. Arrows point towards example invasive cells. Scale bar: 50μm **(c)** Quantification of (a-b) shows no significant change in invasive levels. n≥ 20 animals/ genotype. Scale bar: 50μm. Mann-Whitney U test was performed to determine statistical significance. *p<0.05; **p<0.01; ***p<0.001 ****p<0,0001.

**Figure S4: Identification of differences in gene expression induced by ING1KD through microarray analysis of the MCF7 transcriptome**

**(a)** PCA analysis of genome-wide mRNA expression shows that samples clearly cluster together based on the experimental conditions used: transfection with ING1 siRNA (green), non-targeting siRNA (red) or no treatment (blue). Each dot represents one sample. **(b-d)** Volcano plot of significance level versus fold change in genetic expression from treated and untreated MCF7 cells. Each black dot represents a gene that had no significant change in expression (grey lines represent cut-off points: FC-∈-[-1.5;-1.5], p<0.01 and FDR<0.05). Each red and green dot represents a gene that was downregulated or upregulated, respectively, compared to the control. Differentially expressed genes between ING1KD cells and UNT or NON-T MCF7 cells are shown in (c) and (d) respectively; (b) is the negative control. Two-way ANOVA analysis was performed to determine statistical significance.

## Supplementary Tables and Data

**Table S1: List of differentially expressed genes in response to ING1 knockdown in MCF7 cells**

**Table S2: Biological processes and pathways significantly enriched by ING1KD DEGs and correlated diseases**

**Table S3: Table of genetic and protein interactions between DEGs following ING1KD and AJ components**

**Data S1: Cytoscape network file for interaction map of genes misregulated by ING1KD that affect cell-cell junctions**

## METHODS

### Transgenic Drosophila stocks and crosses

Fly stocks were raised on standard medium at 18°C and crosses on standard medium with yeast at 25 °C. The following stocks were used: yw, neoFRT19A (Chr X; #1744); UAS-p35 (Chr 2; #6298); sna[Sco]/CyO; UAS-iCasper-noGFP-T2A-HO1}VK00005/TM6B (Chr 2,3; #64186) obtained from Bloomington *Drosophila* Stock Center (Indiana, USA); P{GD12222}v27988 (Chr3; #27988); P{GD12222}v27989 (Chr 3; #27989) from VDRC stock centre (Vienna, Austria); Ubi-p63E-shg:GFP (Chr 2, #109007) from KYOTO Stock Center (DGRC).

Ubx-FLP; neoFRT40A/Cyo-GFP; Pnr-GAL4, UAS-GFP:Moe/TM6b

Ubx-FLP; neoFRT40A, *lgl[4]*/Cyo-GFP; Pnr-GAL4, UAS-GFP:Moe/TM6b

w; tub-Gal80, neoFRT40A; MKRS/TM6b

w; IF/Cyo-GFP; Pnr-GAL4, UAS-GFP:Moe/TM6B

Ubx-FLP, neoFRT19A, tub-GAL80; MKRS/TM6B were lab stocks.

We recombined UAS-iCasper-noGFP-T2A-HO1}VK00005 and P{GD12222}v27988 or P{GD12222}v27989 on the 3^rd^ chromosome to generate new stocks.

The following genotypes were imaged: For WT clones: Ubx-FLP/+; neoFRT40A/ tub-GAL80, FRT40A; Pnr-GAl4, UAS-GFP:Moe/MKRS and Ubx-Flp/+; neoFRT40A/tubGAL80, neoFRT40A; PnrGal4, UAS-GFP:Moe/UAS-iCasper

For *lgl*^*4*^ mutants: Ubx-FLP/+; neoFRT40A, *lgl[4]*/tub-GAL80, neoFRT40A; Pnr-GAl4, UAS-GFP:Moe/MKRS and Ubx-Flp/+; neoFRT40A, *lgl[4]*/tub-GAL80, neoFRT40A; PnrGal4, UAS-GFP:Moe/UAS-iCasper

For p35 overexpression: Ubx-FLP/+; neoFRT40A, *lgl[4]*/tub-GAL80, neoFRT40A; Pnr-GAl4, UAS-GFP:Moe/UAS-p35

For CG7379 knockdown: Ubx-FLP/+; neoFRT40A/tub-GAL80, neoFRT40A; Pnr-GAl4, UAS-GFP:Moe/UAS-RNAi and Ubx-Flp/+; neoFRT40A/tub-GAL80, neoFRT40A; PnrGal4, UAS-GFP:Moe/UAS-RNAi, UAS-iCasper and Ubx-Flp, tub-GAL80, neoFRT19A/ neoFRT19A; PnrGal4, UAS-GFP:Moe/UAS-RNAi

For CG7379 KD in *lgl*^*4*^ mutant clones: Ubx-FLP/+; neoFRT40A, *lgl[4]*/tub-GAL80, neoFRT40A; Pnr-GAl4, UAS-GFP:Moe/UAS-RNAi and Ubx-Flp/+; neoFRT40A, *lgl[4]*/tub-GAL80, neoFRT40A; PnrGal4, UAS-GFP:Moe/UAS-RNAi, UAS-iCasper

For E-cadherin overexpression: neoFRT19A/ Ubx-FLP, neoFRT19A, tub-GAL80; +/Ubi-p63E-shg:GFP; Pnr-GAL4, UAS-GFP:Moe/UAS-RNAi

### Cell stocks and maintenance

MDA-MB-231 cells were obtained from Dr Sally Wheatley, U87 cells from Dr Ruman Rahman and MCF7 cells from Dr Anna Grabowska (Faculty of Medicine & Health Sciences, University of Nottingham). Cells were cultured in DMEM (MDA-MB-231 and U87) or RPMI medium without phenol red (MCF7) (Invitrogen) supplemented with 2 mM L-glu (Sigma-Aldrich) and 10% fetal bovine serum (Sigma-Aldrich) and grown in T75 culture flasks at 37 °C in a 5% CO2 atmosphere.

### Dissections, Immunocytochemistry and TdT-Mediated dUTP Nick-End-Labeling

The dorsal thorax of 20-24 hr old pupae was dissected following the protocol described by Jauffred and Bellaiche (Jauffred and Bellaiche, 2012). The tissue was fixed in 4% formaldehyde for 20□min, blocked and permeabilised with PBS containing 0.2% BSA, 5% NGS, 0.1% Triton X-100 followed by primary antibody incubation overnight at 4° C and 1-hour secondary antibody incubation at room temperature. Additionally, a DAPI staining was performed together with the secondary antibody incubation. GeneTex FluoroGel mounting media (GeneTex) was used for mounting. Prior to antibody incubations, MCF7 cells were fixed with 3% PFA for 30 minutes at RT, permeabilised in 0.2% PBS-Triton-X 10mM Glycine and blocked in 3% BSA.

The following primary antibodies were used to label cell-cell junction proteins: mouse anti E cadherin [1:100, abcam (ab1416)]; rat anti-E-Cad [1:100, DSHB (DCAD2)], Mouse anti-Armadillo (1:100, DSHB), rat anti-alpha-Catenin [1:100, DSHB (DCAD1)], Mouse anti-Fasciclin III (1:400, DSHB), mouse anti-Coracle [1:100, DSHB (C615.16)], mouse anti-DiscsLarge [1:100, DSHB (4F3)]; polarity proteins: rat anti-Crumbs (1:1000, gift from E. Knust) rabbit anti-Baz (1:2,000, gift from A. Wodarz), Rabbit anti-PKC zeta (1:50, Santa Cruz); apoptotic cells: rabbit anti-cleaved Drosophila Dcp-1 (Asp216, Cell Signalling Technology). Secondary antibodies from Molecular Probes were Alexa Fluor 488, 546 and 633.

Additionally, the ApopTag® Red In Situ Apoptosis Detection Kit (S7165, Merck Millipore) was also used to label apoptotic cells following Wells and Johnston protocol (Wells and Johnston, 2012). Tissues were incubated as follows: in a pre-cooled ethanol/PBS (2:1) solution, 5 min, −20°C; 10mM Sodium Citrate pH 6.0, 30 min, 70°C; equilibration buffer, 10 min, RT; TdT enzyme (diluted 30% v/v in reaction buffer), 1 hour, 37° C; Stop/Wash solution, 10 min, RT; anti-dig rhodamine (20% antibody, 50% PBSt, 25% blocking solution provided with the kit, 5% NGS), 30 min, RT; DAPI (3 mM), 3 min, RT.

Images from fixed samples were acquired on a Zeiss LSM880 inverted confocal microscope using a x 40/ 1.30 NA oil Ph3 M27 objective, 0.5μm z-sectioning for polarity and junctional proteins and 1μm z-sectioning for samples stained for apoptosis.

### Live imaging

Using double-sided sticky tape animals with the desired genotype were attached to a custom-made slide which has coverslip bridges on the far edges. A window was then cut in the pupal case and a coverslip with a drop of immersion oil was placed on top of the bridges, touching the notum.

The following inverted confocal microscopes and lenses were used for live imaging: Leica SP2 equipped with a × 40/1.25 NA oil lens; Zeiss LSM880C, x 40/ 1.30 NA oil Ph3 M27 lens; Zeiss LSM5 Exciter AxioObserver, Plan-Apochromat x 40/ 1.30 NA oil lens. Samples were excited with a 458nm laser, and signal was collected at 500– 600nm. Z-series were acquired with a 0.35µm pixel size using 1μm z-sectioning. For time-lapses z-series were acquired every 3 minutes for 2 hours.

### Transfection

Transfection media containing DharmaFect1 (Dharmacon) transfection reagent and 25 nM ON-TARGET plus SMARTpool siRNA against ING1 (Dharmacon) was added on 50-60% confluent MCF7 cells and 60-80% confluent MDA-MB-231 or U97 cells. ON-TARGETplus Non-targeting siRNA (Dharmacon) was used as a negative control. Cells were then incubated at 37 °C with 5% CO2 for 48 hours.

### Migration and Invasion assay

MDA-MB-231, U87 or MCF7 cells were plated in 5% BSA or 1% FBS DMEM or RPMI medium respectively, in hanging inserts with 8 µm pore (Millipore). 10% FBS appropriate culture medium was used underneath the inserts. For invasion experiments, the inserts were covered a priori with extracellular matrix (Mark Millipore or Corning). Migrating/ invading cells were detached from the bottom of the inserts using accumax cell detachment solution (Merk Millipore) and stained for DNA content using CyQuant GR dye (Thermo Fisher Scientific). The number of invading cells is direct proportional with the fluorescence intensity as CyQuant dye labels the DNA. The fluorescence intensity was measured using 480/520 nm filters and optimal gain on a BMG fluorescent plate reader.

### RNA extraction

Total cellular RNA was extracted using TRIzol (Thermo Fisher Scientific) following the manufacturer’s instructions.

### cDNA synthesis

cDNA was synthetized using RevertAid First Strand cDNA Synthesis Kit [Thermo Fisher Scientific (k1621)] following the manufacturer’s instructions.

### Quantitative Real-Time PCR

Samples were prepared using IQ SYBR Green qPCR Master Mix (BioRad) and non-universal primers (Thermofisher Scientific, Sigma-Aldrich or Eurofins) designed using Primer3web online tool. For DNA amplification and fluorescence measurements an A C1000 thermal cycler CFX96 RT system was used. Relative transcript levels were calculated relative to expression levels of the house keeping gene GAPDH, using the efficiency calibrated model (Pfaffl, 2001).

#### Primer list

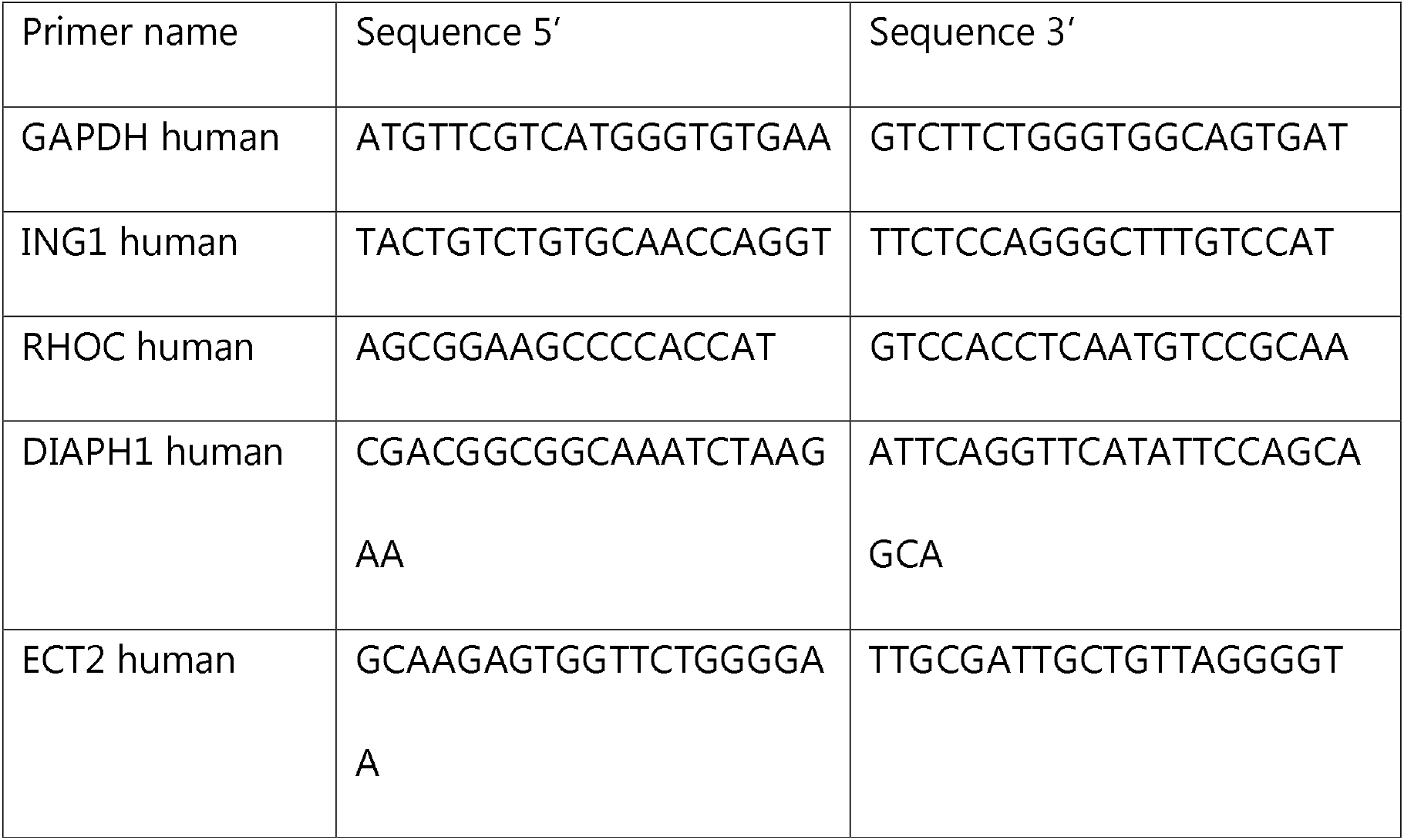

### Whole genome gene expression microarray analysis

Whole-genome transcriptome analysis of transfected and un-transfected (both un-treated and treated with non-targeting siRNA) MCF7 cells was conducted at the Nottingham Arabidopsis Stock Centre (NASC). The RNA concentration and quality was assessed using the Agilent 2100 Bioanalyzer (Agilent Technlogies Inc., Palo Alto, CA) and the RNA 600 Nano Kit (Caliper Life Sciences, Mountain View, CA). Samples with a minimum RNA concentration of 100 ng/μl and RNA Integrity Number (RIN) ≥ 8 were used for gene expression analysis. Single stranded complimentary DNA was prepared from 200 ng of total RNA as per the GeneChip™ WT PLUS Reagent Kit (Applied Biosystems and Affymetrix). Total RNA was first converted to cDNA, followed by *in vivo* transcription to make cRNA. Single stranded cDNA was synthesized, end labelled and hybridized for 16 h at 45°C to Clariom™ S Assay arrays (Thermo Fisher Scientific).

Gene expression data were analysed using Partek Genomics Suite 6.6 software (Partek Incorporated). The raw CEL files were normalized using the RMA background correction with quantile normalization, log base 2 transformation and mean probe-set summarization with adjustment for GC content. Differentially expressed genes (DEGs) were identified by a one-way ANOVA. DEGs were considered significant if p-value with FDR was ≤ 0.05 and fold change of >1.5 or <-1.5.

### GO term enrichment

DEG following ING1 KD were analysed for biological process and pathway enrichment, using Database for Annotation, Visualization and Integrated Discovery (DAVID) Bioinformatics Resources 6.8 (Huang et al., 2007). Functional annotation was performed with EASE score 0.1 and a minimum number of three genes for the corresponding term used as thresholds. Bonferroni correction was applied.

### Interaction map

The interaction network was generated by integrating publicly available data from BioGRID using the GeneMANIA plug-in for the Cytoscape software (Warde-Farley et al., 2010, Shannon et al.). GO molecular function was used as weighting for the ties among nodes; 10 max resultant attributes and 20 max resultant genes were used. The resulting network reflects genetic and physical interactions between these genes.

### Calculations and statistical analysis

Microsoft Excel and Prism (GraphPad) software was used to perform calculations, generate graphs and calculate statistical significance with Student’s t-test, Mann-Whitney U test, Kruskall-Wallis test, and Pearson Correlation test where P>0.05 was considered not significant, *P<0.05, **P<0.01, ***P<0.001, ****P<0.0001 and *r*_*s*_⍰[±0.7, ±1.0] was considered to suggest a strong, *r*_*s*_⍰ [±0.5, ±0.7] a moderate, *r*_*s*_⍰ [±0.3, ±0.5] a weak and *r*_*s*_⍰ [0, ±0.3] a negligible correlation.

For image analysis, standard processing options from Fiji (imageJ) software were used (Schindelin et al., 2012, Cordeliéres).

### Data availability

The accession number for the microarray data reported in this paper is GEO: GSE125438.

