## Supplementary material for "CG7379/ING1 suppress cancer cell invasion by maintaining cell-cell junction integrity": Table S2

**Table S2. Biological processes and pathways significantly enriched by ING1KD DEGs and correlated diseases**

| ID | Term | Bonferroni adjusted p‑value* | No. genes | Genes | |
| --- | --- | --- | --- | --- | --- |
| Biological processes (GOTERM_BP_DIRECT) | | | | | |
| GO:0051301 | Cell division | 2.23E-43 | 97 | ITGB3BP, KIFC1, KNTC1, AURKA, PTTG1, CDCA8, RAD21, CDCA7, MIS18A, CDCA2, CDCA5, CCNA2, CDCA4, CDCA3, STAG1, LIG1, TACC3, NCAPD3, NCAPD2, DCLRE1A, MAD2L1, TIMELESS, RCC2, SPAG5, ZWINT, MCMBP, CDCA7L, NUP43, NEK2, CHEK2, RCC1, SPC25, VRK1, NCAPG2, FBXO5, SKA3, HELLS, ERCC6L, CKAP5, NUF2, ARPP19, SPDL1, CDC20, NDC80, REEP4, FAM64A, SMC1A, BORA, FAM83D, CCNE2, KIF2C, NDE1, MASTL, CDC7, KIF14, CDC6, CDK1, KIF11, DSN1, CCNF, TPX2, PAPD5, UBE2C, CDK2, MCM5, CCND1, CCND3, BUB1B, HAUS8, UBE2S, MAPRE3, CKS1B, HAUS6, NCAPH, NCAPG, BUB1, KLHL42, ZWILCH, BUB3, PSRC1, KIF18B, CENPF, BIRC5, CENPE, CDC25C, KNSTRN, SMC2, CENPJ, SIRT2, CDC25A, SMC4, CCNB1, CCNB2, CKS2, KIF20B, CENPW, MIS18BP1. | |
| Biological processes – continued | | | | | |
| GO:0006260 | DNA replication | 9.46E-37 | 62 | CLSPN, DBF4, NAP1L1, MCM10, CDT1, MCM8, CDC45, MCM7, ORC6, ORC1, CDC7, CDK1, CDC6, DTL, LIG1, POLE, TOPBP1, MCM2, MCM3, RNASEH2A, MCM4, RMI1, MCM5, CDK2, MCM6, RFC5, RFC3, RFC4, TIMELESS, RFC2, RRM2, RRM1, DSCC1, BLM, TICRR, KIAA0101, POLA1, CHEK1, POLA2, RPA3, RPA2, POLE2, FEN1, EXO1, GINS2, SSRP1, GINS3, NASP, GINS4, BRIP1, CDC25C, BRCA1, CDC25A, POLD3, DNA2, POLD4, PCNA, CHTF18, RBM14, CHAF1B, BARD1, DUT. | |
| GO:0007067 | Mitotic nuclear division | 1.13E-31 | 71 | ITGB3BP, KIF22, BORA, KNTC1, PKMYT1, AURKA, PTTG1, AURKB, FAM83D, KIF2C, RAD21, INCENP, MIS18A, CDCA2, MASTL, CDCA5, CCNA2, ASPM, CDCA3, STAG1, CDK1, CDC6, KIF11, KIF15, CCNF, TPX2, PAPD5, PBK, CDK2, DCLRE1A, RCC2, TIMELESS, MCMBP, BUB1B, HAUS8, MAPRE3, NUP43, HAUS6, NEK2, ANLN, CEP55, RCC1, SPC25, VRK1, NCAPG2, BUB1, SKA3, FBXO5, KLHL42, ZWILCH, HELLS, ERCC6L, CENPN, CKAP5, ARPP19, NUF2, CENPF, NDC80, CDC20, BIRC5, CDC25C, REEP4, SIRT2, CDC25A, FAM64A, CCNB2, PLK1, KIF20B, CENPW, MIS18BP1, CIT. | |
| Biological processes – continued | | | | | |
| GO:0007062 | Sister chromatid cohesion | 2.47E-31 | 48 | ITGB3BP, KIF22, RAD51C, KNTC1, RANGAP1, AURKB, SPC25, KIF2C, CDCA8, NDE1, RAD21, DDX11, CENPA, INCENP, BUB1, ZWILCH, CDCA5, BUB3, STAG1, ERCC6L, CENPO, CENPN, CENPM, CKAP5, DSN1, CENPQ, CENPP, NUF2, KIF18A, CENPF, NUP85, NDC80, SPDL1, CENPE, BIRC5, CDC20, CENPK, CENPI, MAD2L1, RCC2, PLK1, ZWINT, MCMBP, BUB1B, CENPU, NUP107, SMC1A, NUP43. | |
| GO:0000082 | G1/S transition of mitotic cell cycle | 2.34E-24 | 42 | CRLF3, DBF4, POLA1, PKMYT1, POLA2, RCC1, MCM10, RPA3, CDT1, CCNE2, PRIM1, RPA2, TYMS, CDC45, MCM8, MCM7, POLE2, CDKN2C, PRIM2, FBXO5, RANBP1, ORC6, ORC1, CDCA5, CDC7, CDK1, CDC6, POLE, SKP2, MCM2, MCM3, CDKN3, MCM4, MCM5, CDK2, CDC25A, MCM6, CCND1, CDKN1A, DHFR, RRM2, PCNA. | |
| GO:0000722 | Telomere maintenance via recombination | 2.97E-17 | 22 | RAD51C, XRCC3, LIG1, POLE, POLA1, BRCA2, POLA2, RAD51, RPA3, RFC5, POLD3, PRIM1, POLD4, RPA2, DNA2, RFC3, RFC4, POLE2, RFC2, PRIM2, PCNA, FEN1. | |
| Biological processes – continued | | | | | |
| GO:0006281 | DNA repair | 4.58E-16 | 51 | KIF22, CLSPN, RAD51C, XRCC3, XRCC2, PTTG1, DDX11, FANCI, FANCA, FANCC, CDK1, GEN1, LIG1, POLE, TOPBP1, CDK2, RAD51, RFC5, UHRF1, RFWD3, FANCD2, RAD18, DMC1, UBE2T, BLM, ENDOV, TICRR, FOXM1, UNG, POLA1, CHEK1, RPA3, POLE2, POLQ, FEN1, ERCC1, EXO1, MSH6, SSRP1, RAD51AP1, MSH2, EME1, RAD54L, BRCA1, TEX15, PARPBP, PARP3, SMC1A, RBM14, PARP2, CHAF1B. | |
| GO:0006270 | DNA replication initiation | 1.06E-15 | 21 | CDC7, CDC6, GINS4, POLE, POLA1, TOPBP1, POLA2, MCM2, MCM10, MCM3, MCM4, MCM5, MCM6, CCNE2, PRIM1, CDC45, MCM7, POLE2, PRIM2, ORC6, ORC1. | |
| GO:0006334 | Nucleosome assembly | 2.47E-15 | 36 | HMGB2, NAP1L1, HIST2H4A, HIST2H4B, HIST1H2BO, HIST1H2BM, HIST1H4A, HIST1H2BL, CENPA, HIST1H2BI, HIST1H2BJ, H2AFX, HIST1H4C, HIST1H4D, ASF1B, HIST3H2BB, H1F0, HIST2H3A, HIST1H1E, HIST1H1D, HIST1H1C, HIST1H2BE, HIST1H2BF, HIST1H1B, NASP, HIST1H2BG, HIST1H2BH, ANP32E, MCM2, HIST2H2BF, HIST1H3A, HIST1H3B, HIST1H3D, HIST1H3F, HIST1H3G, HIST1H3I. | |
| Biological processes – continued | | | | | |
| GO:0000070 | Mitotic sister chromatid segregation | 7.77E-11 | 16 | KIFC1, NEK2, DSN1, NUSAP1, KIF18B, NDC80, ESPL1, KNSTRN, SMC4, MAD2L1, PLK1, CENPA, SPAG5, ZWINT, SMC1A, BUB3. | |
| GO:0000086 | G2/M transition of mitotic cell cycle | 1.19E-10 | 33 | HAUS6, NEK2, FOXM1, BORA, CEP78, PKMYT1, AURKA, CHEK1, CHEK2, HMMR, NDE1, MASTL, TUBB4B, CDK1, WNT10B, CCP110, CKAP5, ARPP19, SKP2, TPX2, BIRC5, CDC25C, CENPJ, CDK2, CDC25A, CCNB1, CDKN1A, PLK4, CCNB2, PLK1, HAUS8, CIT, MELK. | |
| GO:0006271 | DNA strand elongation involved in DNA replication | 1.42E-10 | 13 | GINS1, GINS2, GINS3, GINS4, POLA1, POLA2, PRIM1, POLD3, POLD4, RFC3, RFC4, PRIM2, PCNA. | |
| GO:0000731 | DNA synthesis involved in DNA repair | 1.64E-10 | 18 | EXO1, RAD51C, XRCC3, RAD51AP1, XRCC2, BLM, POLE, POLA1, BRCA2, BRIP1, RMI1, BRCA1, RAD51, POLD3, POLD4, DNA2, RFC3, BARD1. | |
| Biological processes – continued | | | | | |
| GO:0034080 | CENP-A containing nucleosome assembly | 7.5E-10 | 19 | ITGB3BP, CENPO, CENPN, CENPM, CENPQ, CENPP, HIST2H4A, CENPK, CENPI, HIST2H4B, HIST1H4A, HJURP, CENPA, MIS18A, CENPW, HIST1H4C, HIST1H4D, MIS18BP1, CENPU. | |
| GO:0007059 | Chromosome segregation | 8.63E-09 | 22 | CENPN, KIF11, NEK2, DSN1, NUF2, CENPF, NDC80, CENPE, RCC1, KNSTRN, ESCO2, BRCA1, SPC25, NDE1, SPAG5, HJURP, INCENP, MIS18A, CDCA2, CENPW, SKA3, TOP2A. | |
| GO:0006302 | Double-strand break repair | 3.68E-07 | 20 | APLF, MSH2, LIG1, EME1, MND1, BRIP1, PRKDC, BRCA2, CHEK2, ESCO2, BRCA1, RAD21, TDP2, PARP3, H2AFX, POLQ, CDCA5, ERCC1, FEN1, TRIP13. | |
| GO:0000281 | Mitotic cytokinesis | 5.59E-07 | 14 | CKAP2, KIF23, BBS4, KIF4A, CENPA, PLK1, NUSAP1, ANLN, STMN1, CEP55, RACGAP1, SPAST, KIF20A, MYH10. | |
| Biological processes – continued | | | | | |
| GO:0008283 | Cell proliferation | 1.69E-06 | 48 | STIL, CKS1B, CDK5R1, CDV3, ZAK, ERBB2, E2F8, DTYMK, POLA1, PRKDC, AURKB, MCM10, TRAIP, FAM83D, TYMS, KDM1A, KIF2C, MCM7, CSE1L, PRMT5, CKLF, BUB1, ERCC1, TFDP1, POLR3G, CDK1, MKI67, NASP, DLGAP5, KIF15, SKP2, PIM1, TPX2, CENPF, ITGA2, TACC3, CDC25C, CDC25A, IFNAR2, UHRF1, PLK1, CHRM1, PCNA, CKS2, BUB1B, TCF19, MELK, MYH10. | |
| GO:0007080 | Mitotic metaphase plate congression | 1.73E-06 | 15 | KIF14, KIF22, KIFC1, CHMP4C, PSRC1, KIF18A, CENPE, SPDL1, CEP55, CCNB1, KIF2C, CDCA8, CDCA5, KPNB1, PINX1. | |
| GO:0000732 | Strand displacement | 1.85E-06 | 13 | EXO1, RAD51C, DNA2, XRCC3, RAD51AP1, XRCC2, BLM, BRCA2, BRIP1, RMI1, BRCA1, RAD51, BARD1. | |
| GO:0007049 | Cell cycle | 2.48E-06 | 35 | E2F2, CKS1B, DBF4B, USP2, LIN9, FOXM1, DTYMK, AURKA, AURKB, WTAP, DDX12P, CASP8AP2, HJURP, SIK1, ING1, RBL2, NASP, GMNN, RBL1, PIM1, SUV39H1, CDC20, MCM2, CDKN3, ESCO2, BRCA1, SUV39H2, CCND1, UHRF1, CCND3, PTP4A1, CHTF18, SIAH2, CHAF1B, KCTD11. | |
| Biological processes – continued | | | | | |
| GO:0006335 | DNA replication-dependent nucleosome assembly | 2.51E-06 | 14 | NASP, HIST2H4A, HIST2H4B, HIST1H4A, HIST1H3A, HIST1H3B, HIST1H4C, HIST1H3D, HIST1H4D, HIST1H3F, ASF1B, HIST1H3G, CHAF1B, HIST1H3I. | |
| GO:0036297 | Interstrand cross-link repair | 1.27E-05 | 16 | XRCC3, RAD51AP1, USP1, EME1, RAD51, RPA3, RPA2, DCLRE1A, FANCI, FANCD2, FANCE, FANCA, UBE2T, ERCC1, FANCB, FANCC. | |
| GO:0045814 | Negative regulation of gene expression, epigenetic | 1.74E-05 | 16 | HIST2H3A, EZH2, HIST2H4A, HIST2H4B, PHF19, HIST1H4A, HIST1H3A, HIST1H3B, DNMT1, HIST1H4C, HIST1H3D, HIST1H4D, HIST1H3F, DNMT3B, HIST1H3G, HIST1H3I. | |
| GO:0000183 | Chromatin silencing at rDNA | 2.06E-05 | 14 | HIST2H3A, SUV39H1, HIST2H4A, SIRT2, HIST2H4B, HIST1H4A, HIST1H3A, HIST1H3B, HIST1H4C, HIST1H3D, HIST1H4D, HIST1H3F, HIST1H3G, HIST1H3I. | |
| GO:0000724 | Double-strand break repair via homologous recombination | 2.21E-05 | 19 | RAD51C, XRCC3, RAD51AP1, MMS22L, XRCC2, BLM, TONSL, GEN1, BRCA2, RAD54L, BRCA1, RAD51, RPA3, RPA2, MCM8, RAD54B, H2AFX, POLQ, FEN1. | |
| Biological processes – continued | | | | | |
| GO:1901796 | Regulation of signal transduction by p53 class mediator | 5.28E-05 | 24 | EXO1, SSRP1, CDK5R1, BLM, TAF5, TPX2, BRIP1, AURKA, CHEK1, TOPBP1, AURKB, CHEK2, RMI1, BRCA1, CDK2, RPA3, RFC5, RPA2, DNA2, RFC3, RFC4, RFC2, PRMT5, BARD1. | |
| GO:0051726 | Regulation of cell cycle | 5.28E-05 | 24 | E2F2, RBL2, DTL, FIGNL1, FOXM1, LIN9, CCNF, PRR11, RBL1, SKP2, KIAA0101, PKMYT1, CENPF, CDC25C, MYBL2, SIRT2, CDC25A, CCNE2, CCNB1, CCNB2, PLK1, CDK12, MASTL, MED1. | |
| GO:0016925 | Protein sumoylation | 8.28E-05 | 23 | BLM, NUP85, BIRC5, NUP188, RANGAP1, AURKB, NUP155, BRCA1, NDC1, CDCA8, RAD21, AAAS, MDC1, INCENP, NUP205, NUP50, PCNA, NUP107, SMC1A, SCMH1, TOP2A, NUP43, STAG1. | |
| GO:0007018 | Microtubule-based movement | 1.01E-04 | 19 | KIFC2, KIF14, KIF23, KIF22, KIFC1, KIF4A, KIF11, KIF15, KIF18A, DNAH3, KIF18B, CENPE, KIF16B, RACGAP1, KIF2C, KIFAP3, KIF20B, DYNLRB2, KIF20A. | |
| Biological processes – continued | | | | | |
| GO:0000083 | Regulation of transcription involved in G1/S transition of mitotic cell cycle | 1.04E-04 | 11 | CDK1, CDC6, TYMS, CDC45, DHFR, RRM2, PCNA, POLA1, FBXO5, ORC1, CDT1. | |
| GO:0031047 | Gene silencing by RNA | 1.52E-04 | 22 | HIST2H3A, NUP85, NUP188, HIST2H4A, NUP155, HIST2H4B, NDC1, AAAS, HIST1H4A, NUP205, NUP50, HIST1H3A, AGO1, HIST1H3B, HIST1H3D, HIST1H4C, HIST1H4D, NUP107, HIST1H3F, NUP43, HIST1H3G, HIST1H3I. | |
| GO:0042769 | DNA damage response, detection of DNA damage | 1.54E-04 | 13 | DTL, USP1, RPA3, POLD3, RFC5, POLD4, RPA2, RFC3, RFC4, RFC2, PCNA, DNAJA1, RAD18. | |
| GO:0006297 | Nucleotide-excision repair, DNA gap filling | 1.71E-04 | 11 | RFC5, POLD3, POLD4, RPA2, RFC3, RFC4, RFC2, LIG1, POLE, PCNA, RPA3. | |
| GO:0007077 | Mitotic nuclear envelope disassembly | 2.20E-04 | 14 | CDK1, NUP85, NUP188, NUP155, CCNB1, NDC1, VRK1, AAAS, CCNB2, PLK1, NUP205, NUP50, NUP107, NUP43. | |
| Biological processes – continued | | | | | |
| GO:0032200 | Telomere organization | 6.44E-02 | 11 | HIST1H4A, HIST1H3A, HIST1H3B, HIST1H3D, HIST1H4C, HIST1H4D, HIST1H3F, HIST2H4A, HIST1H3G, HIST1H3I, HIST2H4B. | |
| GO:0007051 | Spindle organization | 6.56E-04 | 9 | KIF11, CKAP5, SPAG5, TTK, AURKA, RANBP1, AURKB, KNSTRN, ASPM. | |
| GO:0051290 | Protein heterotetramerization | 0.0010 | 13 | HIST2H4A, HIST2H4B, HIST1H4A, RRM2, HIST1H3A, RRM1, HIST1H3B, HIST1H4C, HIST1H3D, HIST1H4D, HIST1H3F, HIST1H3G, HIST1H3I. | |
| GO:0019985 | Translesion synthesis | 0.0012 | 12 | RFC5, POLD3, POLD4, RPA2, RFC3, RFC4, DTL, RFC2, PCNA, KIAA0101, RPA3, REV3L. | |
| GO:0007052 | Mitotic spindle organization | 0.0020 | 11 | CCNB1, STIL, SPC25, KIF11, GPSM2, TTK, NDC80, AURKA, STMN1, RCC1, SMC1A. | |
| GO:0007131 | Reciprocal meiotic recombination | 0.0020 | 11 | RAD51C, MSH6, XRCC3, RAD21, XRCC2, MSH2, MND1, RAD54B, DMC1, RAD51, TRIP13. | |
| Biological processes – continued | | | | | |
| GO:0006977 | DNA damage response, signal transduction by p53 class mediator resulting in cell cycle arrest | 0.0027 | 15 | E2F1, CDK1, RBL2, E2F7, AURKA, CHEK2, CDC25C, CENPJ, CNOT6, CDK2, GTSE1, CCNB1, CDKN1A, PCNA, TFDP1. | |
| GO:0042276 | Error-prone translesion synthesis | 0.0034 | 9 | RFC5, RPA2, RFC3, RFC4, POLE2, RFC2, PCNA, RPA3, REV3L. | |
| GO:0000910 | Cytokinesis | 0.0050 | 13 | KIF23, PRC1, BRCA2, ESPL1, BIRC5, ECT2, ARL3, PLK1, INCENP, RHOC, CIT, NEK7, KIF20A. | |
| GO:0071897 | DNA biosynthetic process | 0.0051 | 10 | TYMS, LIN9, LIG1, PRIM2, CENPF, PAPD5, POLA2, POLQ, TERT, TK1. | |
| Biological processes – continued | | | | | |
| GO:0006974 | Cellular response to DNA damage stimulus | 0.0062 | 28 | XRCC3, BLM, APLF, KIAA0101, CHEK1, CHEK2, MCM10, MCM8, MCM7, DDX11, WDR76, H2AFX, MASTL, POLQ, TOP2A, DTL, SUV39H1, TOPBP1, ATAD5, BRCA1, RAD51, CDKN1A, CCND1, TIMELESS, RAD18, UBE2T, UBE2E2, BARD1. | |
| GO:0007076 | Mitotic chromosome condensation | 0.0069 | 8 | NCAPH, NCAPG, NUSAP1, CDCA5, SMC2, NCAPD3, SMC4, NCAPD2. | |
| GO:0006284 | Base-excision repair | 0.0098 | 11 | HMGB1, RPA2, MPG, DNA2, LIG1, UNG, NEIL3, POLQ, PARP2, FEN1, RPA3. | |
| GO:0060968 | Regulation of gene silencing | 0.0119 | 7 | HIST1H3A, HIST1H3B, HIST1H3D, HIST1H3F, HIST1H3G, CDK2, HIST1H3I. | |
| GO:1900264 | Positive regulation of DNA-directed DNA polymerase activity | 0.0129 | 6 | RFC5, RFC3, RFC4, RFC2, CHTF18, DSCC1. | |
| GO:0032508 | DNA duplex unwinding | 0.0135 | 12 | GINS1, GINS2, DNA2, CDC45, BLM, DDX11, GINS4, BRIP1, RAD54B, MCM3, MCM5, DDX12P. | |
| Biological processes – continued | | | | | |
| GO:0045815 | Positive regulation of gene expression, epigenetic | 0.0157 | 14 | HIST2H3A, KAT2B, DEK, HIST2H4A, HIST2H4B, HIST1H4A, HIST1H3A, HIST1H3B, HIST1H4C, HIST1H3D, HIST1H4D, HIST1H3F, HIST1H3G, HIST1H3I. | |
| GO:0006342 | Chromatin silencing | 0.0171 | 12 | HIST1H2AB, HIST2H2AA3, HIST2H2AB, HIST1H2AG, HIST1H2AE, H2AFZ, HIST1H2AI, H2AFX, HIST1H2AJ, HIST1H2AM, HIST1H2AL, SIRT2. | |
| GO:0006310 | DNA recombination | 0.0213 | 16 | EXO1, RAD51C, HMGB1, XRCC3, HMGB3, BLM, LIG1, RAD54L, BRCA1, RAD51, RPA3, RAD21, PSMC3IP, RBM14, RBPJ, ERCC1. | |
| GO:0051310 | Metaphase plate congression | 0.0228 | 7 | FAM83D, KIF22, KIF2C, CENPQ, CENPF, CENPE, NDC80. | |
| GO:0007088 | Regulation of mitotic nuclear division | 0.0263 | 9 | MKI67, NEK2, PRMT5, BORA, KIF20B, FBXO5, PKMYT1, RCC1, CDC25C. | |
| Biological processes – continued | | | | | |
| GO:0000079 | Regulation of cyclin-dependent protein serine/threonine kinase activity | 0.0284 | 11 | CCNE2, CDC6, CDK5R1, CDKN1A, BLM, CDKN2C, PKMYT1, CDC25C, CDKN3, CCNA2, CDC25A. | |
| GO:0006312 | Mitotic recombination | 0.0403 | 7 | RAD51C, XRCC2, RAD54B, DMC1, TOP2A, ERCC1, RAD51. | |
| GO:0010212 | Response to ionizing radiation | 0.0408 | 12 | RAD51C, XRCC3, RFWD3, XRCC2, TICRR, RRM1, H2AFX, TOPBP1, DMC1, DNMT3B, RAD54L, BRCA1. | |
| Pathways (KEGG) | | | | | |
| hsa04110 | Cell cycle | 7.34E-26 | 47 | | E2F1, E2F2, DBF4, PKMYT1, TTK, PRKDC, CHEK1, CHEK2, PTTG1, CCNE2, CDC45, RAD21, MCM7, CDKN2C, BUB1, ORC6, CCNA2, ORC1, BUB3, TFDP1, STAG1, CDC7, CDK1, CDC6, RBL2, RBL1, SKP2, ESPL1, CDC20, MCM2, MCM3, CDC25C, MCM4, MCM5, CDK2, CDC25A, MCM6, CCNB1, CCND1, CDKN1A, CCNB2, MAD2L1, CCND3, PLK1, PCNA, BUB1B, SMC1A. |
| Pathways – continued | | | | | |
| hsa03030 | DNA replication | 1.19E-20 | 25 | | POLA1, POLA2, RPA3, PRIM1, RPA2, MCM7, POLE2, PRIM2, FEN1, LIG1, POLE, MCM2, MCM3, RNASEH2A, MCM4, MCM5, MCM6, POLD3, RFC5, POLD4, DNA2, RFC3, RFC4, RFC2, PCNA. |
| hsa05322 | Systemic lupus erythematosus | 1.25E-14 | 37 | HLA-DQB1, HIST1H2AB, HIST2H2AA3, HLA-DRB1, HIST1H2AG, HIST1H2AE, HIST2H4A, HIST2H4B, HIST1H2BO, HIST2H2AB, HIST1H2BM, HIST1H4A, HIST1H2BL, HIST1H2BI, HIST1H2BJ, H2AFZ, H2AFX, HIST1H4C, HIST1H4D, HIST3H2BB, HIST2H3A, HIST1H2BE, HIST1H2BF, HIST1H2BG, HIST1H2BH, HIST2H2BF, HIST1H3A, HIST1H3B, SNRPB, HIST1H2AI, HIST1H3D, HIST1H2AJ, HIST1H3F, HIST1H2AM, HIST1H2AL, HIST1H3G, HIST1H3I. | |
| hsa03460 | Fanconi anemia pathway | 1.39E-09 | 20 | RAD51C, BLM, USP1, EME1, BRCA2, BRIP1, RMI1, BRCA1, RAD51, RPA3, RPA2, FANCD2, FANCI, FANCE, FANCA, UBE2T, ERCC1, FANCB, FANCC, REV3L. | |
| Pathways – continued | | | | | |
| hsa05034 | Alcoholism | 1.87E-08 | 34 | HIST1H2AB, HIST2H2AA3, HIST1H2AG, HIST1H2AE, HIST2H4A, HIST2H4B, HIST1H2BO, HIST2H2AB, HIST1H2BM, HIST1H4A, HIST1H2BL, HIST1H2BI, HIST1H2BJ, H2AFZ, H2AFX, HIST1H4C, HIST1H4D, HIST3H2BB, HIST2H3A, HIST1H2BE, HIST1H2BF, HIST1H2BG, HIST1H2BH, HIST2H2BF, HIST1H3A, HIST1H3B, HIST1H2AI, HIST1H3D, HIST1H2AJ, HIST1H3F, HIST1H2AM, HIST1H2AL, HIST1H3G, HIST1H3I. | |
| hsa03430 | Mismatch repair | 6.17E-08 | 13 | EXO1, MSH6, MSH2, LIG1, RPA3, POLD3, RFC5, RPA2, POLD4, RFC3, RFC4, RFC2, PCNA. | |
| hsa03440 | Homologous recombination | 1.78E-06 | 13 | RAD51C, XRCC3, XRCC2, BLM, EME1, BRCA2, RAD54L, RAD51, RPA3, POLD3, RPA2, POLD4, RAD54B. | |
| hsa05203 | Viral carcinogenesis | 3.71E-06 | 33 | CHEK1, HIST2H4A, HIST2H4B, HIST1H2BO, CCNE2, HIST1H2BM, HIST1H4A, GSN, HIST1H2BL, HIST1H2BI, HIST1H2BJ, RANBP1, HIST1H4C, HIST1H4D, HIST3H2BB, CCNA2, CDK1, KAT2B, RBL2, HIST1H2BE, HIST1H2BF, HIST1H2BG, HIST1H2BH, RBL1, SKP2, CDC20, CDK2, CDKN1A, CCND1, CCND3, HIST2H2BF, SCIN, RBPJ. | |
| Pathways – continued | | | | | |
| hsa03410 | Base excision repair | 1.01E-05 | 13 | HMGB1, LIG1, NEIL3, UNG, POLE, POLD3, POLD4, MPG, POLE2, PCNA, PARP3, PARP2, FEN1. | |
| hsa00240 | Pyrimidine metabolism | 1.90E-04 | 20 | POLR3G, POLE, DTYMK, POLA1, DCK, CTPS1, POLA2, PNP, TK1, PRIM1, POLD3, POLD4, TYMS, POLE2, RRM2, RRM1, PRIM2, UCK2, ENTPD1, DUT. | |
| hsa03420 | Nucleotide excision repair | 7.64E-04 | 13 | LIG1, POLE, RPA3, POLD3, RFC5, POLD4, RPA2, RFC3, RFC4, POLE2, RFC2, PCNA, ERCC1. | |
| hsa04115 | P53 signalling pathway | 0.0075 | 14 | CDK1, CHEK1, CHEK2, CDK2, GTSE1, CCNE2, CCNB1, EI24, CCND1, CDKN1A, CCNB2, CCND3, CD82, RRM2. | |
| hsa04114 | Oocyte meiosis | 0.0121 | 18 | CDK1, ADCY6, PKMYT1, AURKA, CDC20, ESPL1, PTTG1, CDC25C, CDK2, CCNB1, CCNE2, IGF1R, CCNB2, MAD2L1, PLK1, BUB1, FBXO5, SMC1A. | |
| Diseases (GAD) | | | | | |
|  | Breast cancer | 1.28E-27 | 104 | RAD51C, XRCC3, RNASEL, XRCC2, TTK, AURKA, PTTG1, AURKB, CXCL12, CDKN2C, H2AFX, ASPM, LIG1, POLE, SKP2, ESPL1, TOPBP1, TACC3, NCAPD2, MAD2L1, SPAG5, SNRPB, RAD18, NEK7, BBS4, HLA-DRB1, BLM, NEK2, ERBB2, RABGAP1L, CHEK1, CHEK2, RHOBTB2, DNMT3B, CKAP5, EME1, NUF2, BRIP1, PPFIBP2, BRCA2, ITGA2, CDC20, NDC80, RAD54L, BRCA1, CDKN1A, PLK4, DHFR, PLK1, PCNA, TUBD1, KIF23, CLSPN, CDT1, MTHFD1, KIF2C, MCM7, FANCE, RANBP1, FANCA, TOP2A, TERT, FANCB, FANCC, CDK1, KIF11, GEN1, TPX2, SPINT1, MCM2, MCM3, MCM4, MCM5, CDK2, MCM6, RAD51, CCND1, CCND3, FANCD2, BUB1B, DMC1, MED1, HLA-DQB1, CALCR, YPEL5, PRKDC, RPA3, IGF1R, TYMS, RPA2, BUB1, GPSM2, FEN1, ERCC1, BUB3, PINX1, MSH6, SHMT1, MSH2, GMNN, CENPE, CDC25C, CCNB1, BARD1. | |
| Diseases – continued | | | | | |
|  | Bladder cancer | 5.42E-08 | 64 | RAD51C, XRCC3, RNASEL, XRCC2, AURKA, H2AFX, FANCA, IFNGR2, CCNA2, TERT, FANCC, RET, LIG1, POLE, PIM1, TACC3, RAD51, RFC5, IFNAR2, CCND1, RFC3, RFC4, CCND3, FANCD2, RFC2, PSCA, REV3L, CALCR, BLM, ERBB2, UNG, PRKDC, CHEK1, CHEK2, POLA2, MYBL2, RPA3, TYMS, IGF1R, RPA2, POLE2, POLQ, ERCC1, EXO1, SHMT1, MSH6, MSH2, GGH, EPHX2, BRIP1, BRCA2, STAT1, CDC25C, RAD54L, BRCA1, CDC25A, POLD3, MPG, CDKN1A, DHFR, PLK1, PCNA, PARP3, BARD1. | |
|  | Pancreatic neoplasms | 4.63E-07 | 18 | EXO1, MSH6, MSH2, USP1, BRCA2, BRIP1, CHEK1, TOPBP1, BRCA1, RAD51, CDKN1A, MDC1, FANCD2, FANCE, RAD54B, FANCA, ABCC5, FANCC. | |
|  | Ovarian cancer | 8.14E-07 | 49 | SAT1, E2F1, RAD51C, E2F2, XRCC3, XRCC2, AURKA, PTTG1, MTHFD1, CCNE2, SPRY1, PLOD2, CDKN2C, CCNA2, TERT, H1F0, WNT10B, RBL2, RBL1, SKP2, MCM2, CDK2, RAD51, MCM6, CCND1, CCND3, HLA-DQB1, HLA-DRB1, CRLF3, ERBB2, CHEK2, TYMS, BUB1, ERCC1, TFDP1, MSH6, MKI67, MSH2, BRCA2, CDC20, STAT1, CDC25A, BRCA1, CCNB1, CDKN1A, CCNB2, ID2, PLK1, PTP4A1. | |
| Diseases – continued | | | | | |
|  | Chronic renal failure | 9.06E-06 | 85 | E2F2, XRCC3, XRCC2, CYP2S1, DBF4, E2F7, PINK1, CXCL12, TTLL12, GSTM2, SLC16A1, GSTM4, FANCE, FANCA, CCNA2, IFNGR2, CDC6, LIG1, ADGRE5, NEIL3, POLE, PBK, TACC3, MCM5, CDK2, NCAPD2, DCLRE1A, CCND1, RFC3, CCND3, SBF2, RRM1, NUCB2, RAD18, BUB1B, DMC1, REV3L, DCBLD2, CCL3, PPP2R3A, HLA-DRB1, BLM, TICRR, NEK2, ERBB2, FOXM1, DCK, CHEK1, POLA2, CHEK2, HMMR, SLC29A1, TYMS, IGF1R, FMO5, PTK2B, DNAJA1, CERK, BCAS3, POLQ, ERCC1, EXO1, MSH6, HERPUD1, MSH2, EME1, BRCA2, ITGA2, NDC80, CDC20, BIRC5, LHPP, BRCA1, CYP4B1, IKBKE, CDKN1A, PLK4, SYNE2, AOX1, PCNA, PHGDH, PARP3, RAD54B, PARP2, ABCC5. | |
| Diseases – continued | | | | | |
|  | Leukemia, Lymphocytic, Chronic, B-Cell | 7.01E-05 | 32 | HLA-DQB1, E2F2, XRCC2, HLA-DRB1, EZH2, PRKDC, AURKB, SP110, CXCL12, CDC45, RAD21, MDC1, TOP2A, EXO1, CPT1B, CDC6, RBL2, MSH2, GMNN, POLE, EPHX2, CENPF, BIRC5, CDC25C, MCM4, ESCO2, MCM5, CYP4B1, RAD51, CCND1, BUB1B, REV3L. | |
|  | Esophageal adenocarcinoma | 1.12E-04 | 43 | XRCC3, XRCC2, ERBB2, CHEK1, AURKA, CHEK2, MTHFD1, SLC29A1, TYMS, IGF1R, CASP7, FANCA, CCNA2, IFNGR2, DNMT3B, TERT, ERCC1, EXO1, MSH6, SHMT1, RET, MSH2, LIG1, RBL1, GGH, BRCA2, ITGA2, BIRC5, CDC25C, BRCA1, CDC25A, RAD51, CCNB1, IFNAR2, CCND1, CDKN1A, CCND3, RRM1, PCNA, RAD18, BUB1B, BARD1, REV3L. | |
|  | Glioma | 1.37E-04 | 17 | HLA-DQB1, PHLDB1, XRCC3, XRCC2, HLA-DRB1, GEN1, LIG1, ERBB2, EME1, BRCA2, PRKDC, RAD54B, RAD54L, BRCA1, TERT, ERCC1, RAD51. | |
| Diseases – continued | | | | | |
|  | Lung cancer | 2.09E-04 | 54 | FHIT, XRCC3, RNASEL, XRCC2, EZH2, AURKA, CASP7, IFNGR2, CCNA2, FANCA, TERT, CDK1, RET, C5AR1, LIG1, PIM1, RAD51, IFNAR2, CCND1, CCND3, RRM1, BUB1B, RAD18, HLA-DQB1, CALCR, BLM, HLA-DRB1, ERBB2, PRKDC, CHEK1, CHEK2, MYBL2, TYMS, IGF1R, DNMT3B, FEN1, ERCC1, EXO1, MSH6, MSH2, EPHX2, GGH, BRCA2, BRIP1, CDC25C, STAT1, RAD54L, BRCA1, CDC25A, CDKN1A, DHFR, PLK1, PCNA, BARD1. | |
|  | Hematologic Neoplasms | 0.0142 | 12 | RFC5, HLA-DQB1, MPG, RFC3, RFC4, HLA-DRB1, RFC2, LIG1, UNG, POLE, PCNA, FEN1. | |
|  | Colorectal Neoplasms | 0.0168 | 18 | MSH6, XRCC3, XRCC2, MSH2, LIG1, BRCA2, CHEK2, BRCA1, RAD51, TYMS, CCND1, CDKN1A, CASP8AP2, FANCD2, PCNA, RAD54B, ERCC1, BARD1. | |
|  | Cervical intraepithelial neoplasia grade‑3 | 0.0337 | 12 | EXO1, MSH6, XRCC3, PCNA, BRCA2, BRIP1, RAD54B, STAT1, FANCA, IFNGR2, BRCA1, TERT. | |

*DAVID functional annotation performed on the 06^th^ of June 2020
