## Supplementary material for "CG7379/ING1 suppress cancer cell invasion by maintaining cell-cell junction integrity": Table S3

### Table S3. Table of genetic and protein interactions between DEGs following ING1KD and AJ components

| Interaction | Normalised  maxim weight | Type of interaction |
| --- | --- | --- |
| Genetic interactions |  |  |
| ELF3 - ERBB2 | 0.0050 | DEG |
| TCF7L2 - CTNNB1 | 0.0019 | linker gene; AJ component |
| LMNA - JUP | 3.86E-06 | linker genes |
| CCNA2 - AFDN | 2.48E-06 | AJ components |
| EPHB3 - GPSM2 | 2.21E-06 | DEG; linker genes |
| E2F2 - RHOC | 2.19E-06 | DEG |
| PVR - CCNA2 | 2.01E-06 | DEG; linker genes |
| CDH1 - GPSM2 | 1.75E-06 | DEG; AJ component |
| EPHB3 - DIAPH1 | 1.69E-06 | DEG; linker genes |
| CCNB2 - CTNNB1 | 1.43E-06 | DEG; AJ component |
| JUP - E2F2 | 1.41E-06 | DEG; linker genes |
| EPS15 - GPNMB | 1.39E-06 | DEG; linker genes |
| ARHGAP19 - NECTIN1 | 1.37E-06 | DEG; AJ component |
| ARHGAP21 - GPNMB | 1.34E-06 | DEG |
| JUP - CDH1 | 1.27E-06 | linker gene; AJ component |
| PTRM - RHOC | 1.27E-06 | DEG; linker genes |
| CDH1 - PCDHB16 | 1.26E-06 | DEG; AJ component |
| LMNB1 - RACGEF2 | 1.21E-06 | DEG |
| EGFR - DIAPH1 | 1.19E-06 | DEG; linker genes |
| ERBB2 - RACGAP1 | 1.18E-06 | DEG |
| CTNNA1 - CCNA2 | 1.17E-06 | DEG; AJ component |
| TCF7L2 - CCNB2 | 1.14E-06 | DEG; linker genes |
| ADD3 - PCDHB16 | 1.11E-06 | DEG |
| CTNNA3 - GPNMB | 1.06E-06 | DEG; AJ component |
| CCNB2 - CTNNA3 | 1.05E-06 | DEG; AJ component |
| Genetic interactions – continued | | |
| AFF3 - PCDHB16 | 1.03E-06 | DEG |
| LMNB1 - CTNNA2 | 9.74E-07 | DEG; AJ component |
| STON2 - CTNNA2 | 9.42E-07 | DEG; AJ component |
| CCNA2 - ARHGAP21 | 9.24E-07 | DEG |
| EGFR - ERBB2 | 8.94E-07 | DEG; linker genes |
| CDH1 - RBL1 | 8.67E-07 | DEG; AJ component |
| PTRM - DIAPH1 | 8.65E-07 | DEG; linker genes |
| AGO1 - NECTIN1 | 8.44E-07 | DEG; AJ component |
| KDM5B - ECT2 | 8.13E-07 | DEG |
| PTRM - CCNA2 | 8.12E-07 | DEG; linker genes |
| PTRM - PCDHB16 | 8.09E-07 | DEG; linker genes |
| ERBB2 - CTNNB1 | 8.02E-07 | DEG; AJ component |
| CDH1 - NECTIN2 | 7.74E-07 | AJ components |
| RBL1 - ADD3 | 7.67E-07 | DEG |
| CTNNA2 - PCDHB16 | 7.37E-07 | DEG; AJ component |
| KIF23 - MYH10 | 7.31E-07 | DEG |
| AFDN - ADD3 | 6.96E-07 | DEG; AJ component |
| EPHB3 - KDM5B | 6.94E-07 | DEG; linker genes |
| CTNNB1 - RBL1 | 6.89E-07 | DEG; AJ component |
| NECTIN2 - ADD3 | 6.86E-07 | DEG; AJ component |
| EGFR - NECTIN2 | 6.86E-07 | linker gene; AJ component |
| ECT2 - EXOC5 | 6.54E-07 | DEG |
| RBL1 - ARHGAP21 | 6.35E-07 | DEG |
| CTNNB1 - NECTIN2 | 6.16E-07 | AJ components |
| KDM5B - E2F2 | 6.12E-07 | DEG |
| KDM5B - EXOC5 | 6.11E-07 | DEG |
| PVR - MYH10 | 6.05E-07 | DEG; linker genes |
| EPS15 - NECTIN2 | 5.90E-07 | linker gene; AJ component |
| Genetic interactions – continued | | |
| CDH1 - ECT2 | 5.90E-07 | DEG; AJ component |
| ELF3 - RACGEF2 | 5.87E-07 | DEG; linker genes |
| AFDN - RACGEF2 | 5.73E-07 | DEG; AJ component |
| EPHB3 - EXOC5 | 5.58E-07 | DEG; linker genes |
| SP1 - CDH1 | 5.38E-07 | linker gene; AJ component |
| EGFR - ECT2 | 5.22E-07 | DEG; linker genes |
| MYH10 - KDM5B | 5.22E-07 | DEG |
| CTNNA2 - ARHGAP19 | 5.16E-07 | DEG; AJ component |
| CTNNA1 - KDM5B | 5.12E-07 | DEG; AJ component |
| ARHGAP19 - CTNNA3 | 5.10E-07 | DEG; AJ component |
| PVR - CTNNB1 | 5.08E-07 | linker gene; AJ component |
| EPHB3 - CDH1 | 5.04E-07 | linker gene; AJ component |
| SP1 - CTNNA1 | 4.99E-07 | linker gene; AJ component |
| TCF7L2 - NECTIN2 | 4.88E-07 | linker gene; AJ component |
| EGFR - KDM5B | 4.88E-07 | DEG; linker genes |
| ECT2 - AFF3 | 4.82E-07 | DEG |
| EPS15 - NECTIN1 | 4.74E-07 | linker gene; AJ component |
| ARHGAP21 - NECTIN1 | 4.56E-07 | DEG; AJ component |
| EPHB3 - EGFR | 4.46E-07 | linker genes |
| CDH1 - E2F2 | 4.44E-07 | DEG; AJ component |
| ECT2 - ARHGAP21 | 4.32E-07 | DEG |
| EGFR - RACGAP1 | 4.14E-07 | DEG; linker genes |
| EPHB3 - AFF3 | 4.12E-07 | DEG; linker genes |
| KDM5B - RACGEF2 | 4.02E-07 | DEG |
| PTRM - NECTIN1 | 4.01E-07 | linker gene; AJ component |
| EGFR - E2F2 | 3.93E-07 | DEG; linker genes |
| PTRM - ECT2 | 3.80E-07 | DEG; linker genes |
| TCF7L2 - ECT2 | 3.72E-07 | DEG; linker genes |
| Genetic interactions – continued | | |
| PVR - CTNNA3 | 3.71E-07 | linker gene; AJ component |
| EPHB3 - ARHGAP21 | 3.69E-07 | DEG; linker genes |
| CTNNA2 - NECTIN1 | 3.65E-07 | AJ components |
| E2F2 - AFF3 | 3.63E-07 | DEG |
| EPS15 - RACGAP1 | 3.56E-07 | DEG; linker genes |
| PTRM - KDM5B | 3.55E-07 | DEG; linker genes |
| EGFR - CDH1 | 3.54E-07 | linker gene; AJ component |
| CTNNB1 - EXOC5 | 3.52E-07 | DEG; AJ component |
| CTNNA1 - MYH10 | 3.52E-07 | DEG; AJ component |
| SP1 - PTRM | 3.46E-07 | linker genes |
| RACGAP1 - ARHGAP21 | 3.43E-07 | DEG |
| NECTIN3 - ARHGAP21 | 3.40E-07 | DEG; AJ component |
| RACGEF2 - NECTIN3 | 3.39E-07 | DEG; AJ component |
| EPS15 - E2F2 | 3.38E-07 | DEG; linker genes |
| MYH10 - ADD3 | 3.35E-07 | DEG |
| E2F2 - ARHGAP21 | 3.25E-07 | DEG |
| RACGEF2 - EXOC5 | 3.23E-07 | DEG |
| KDM5B - CTNNA3 | 3.20E-07 | DEG; AJ component |
| CTNNB1 - CDH1 | 3.18E-07 | AJ components |
| SP1 - CTNNA2 | 3.15E-07 | linker gene; AJ component |
| EGFR - ADD3 | 3.13E-07 | DEG; linker genes |
| EPS15 - CDH1 | 3.04E-07 | linker gene; AJ component |
| PTRM - NECTIN3 | 2.99E-07 | linker gene; AJ component |
| EPHB3 - CTNNA2 | 2.96E-07 | linker gene; AJ component |
| EGFR - AFF3 | 2.89E-07 | DEG; linker genes |
| EPS15 - MYH10 | 2.88E-07 | DEG; linker genes |
| PTRM - E2F2 | 2.86E-07 | DEG; linker genes |
| EPS15 - CTNNA1 | 2.83E-07 | linker gene; AJ component |
| Genetic interactions – continued | | |
| TCF7L2 - EXOC5 | 2.79E-07 | DEG; linker genes |
| MYH10 - ARHGAP21 | 2.77E-07 | DEG |
| CTNNA1 - ARHGAP21 | 2.72E-07 | DEG; AJ component |
| CTNNA1 - RACGEF2 | 2.71E-07 | DEG; AJ component |
| NECTIN3 - CTNNA3 | 2.70E-07 | AJ components |
| EGFR - ARHGAP21 | 2.59E-07 | DEG; linker genes |
| RACGEF2 - ADD3 | 2.58E-07 | DEG |
| EXOC5 - CTNNA3 | 2.57E-07 | DEG; AJ component |
| EPS15 - AFF3 | 2.49E-07 | DEG; linker genes |
| EPS15 - CTNNB1 | 2.42E-07 | linker gene; AJ component |
| ARHGAP21 - AFF3 | 2.40E-07 | DEG |
| PTRM - CTNNA1 | 2.39E-07 | linker gene; AJ component |
| TCF7L2 - CTNNA1 | 2.34E-07 | linker gene; AJ component |
| CDH1 - CTNNA3 | 2.32E-07 | AJ components |
| PTRM - EGFR | 2.28E-07 | linker genes |
| EPS15 - ARHGAP21 | 2.23E-07 | DEG; linker genes |
| EPS15 - RACGEF2 | 2.22E-07 | DEG; linker genes |
| MYH10 - CTNNA2 | 2.22E-07 | DEG; AJ component |
| CTNNA2 - ADD3 | 2.08E-07 | DEG; AJ component |
| TCF7L2 - EPS15 | 1.92E-07 | linker genes |
| AFF3 - CTNNA3 | 1.90E-07 | DEG; AJ component |
| PTRM - RACGEF2 | 1.88E-07 | DEG; linker genes |
| TCF7L2 - RACGEF2 | 1.84E-07 | DEG; linker genes |
| EPS15 - CTNNA2 | 1.79E-07 | linker gene; AJ component |
| EPS15 - CTNNA3 | 1.77E-07 | linker gene; AJ component |
| ARHGAP21 - CTNNA3 | 1.70E-07 | DEG; AJ component |
| RACGEF2 - CTNNA3 | 1.69E-07 | DEG; AJ component |
| TCF7L2 - PTRM | 1.62E-07 | linker genes |
| Genetic interactions – continued | | |
| PTRM - CTNNA2 | 1.51E-07 | linker gene; AJ component |
| PTRM - CTNNA3 | 1.50E-07 | linker gene; AJ component |
| TCF7L2 - CTNNA2 | 1.48E-07 | linker gene; AJ component |
| CTNNA2 - CTNNA3 | 1.36E-07 | AJ components |
| Protein interactions |  |  |
| CKS1B - CDK1 | 0.0045 | DEG |
| EPS15 - STON2 | 0.0044 | DEG; linker genes |
| KIF23 - RACGAP1 | 0.0039 | DEG |
| EPHB3 - AFDN | 0.0035 | linker gene; AJ component |
| NECTIN3 - NECTIN2 | 0.0031 | AJ components |
| PTRM - CDH1 | 0.0031 | linker gene; AJ component |
| KIF23 - ECT2 | 0.0030 | DEG |
| CKS1B - CDH1 | 0.0028 | DEG; AJ component |
| CCNB2 - CDK2 | 0.0028 | DEG |
| LMNA - LMNB1 | 0.0027 | DEG; linker genes |
| EGFR - CTNND1 | 0.0027 | DEG; linker genes |
| EGFR - CDH1 | 0.0027 | linker gene; AJ component |
| EGFR - CDK1 | 0.0027 | DEG; linker genes |
| EGFR - JUP | 0.0027 | linker genes |
| EGFR - WASF3 | 0.0027 | DEG; linker genes |
| CTNNA1 - CDH1 | 0.0025 | AJ components |
| CDH1 - CTNND1 | 0.0025 | DEG; AJ component |
| CCNB1 - CDK1 | 0.0024 | DEG |
| CCNB2 - CDK1 | 0.0024 | DEG |
| SP1 - E2F2 | 0.0024 | DEG; linker genes |
| RACGAP1 - ECT2 | 0.0021 | DEG |
| PVR - NECTIN1 | 0.0021 | linker gene; AJ component |
| TCF7L2 - CTNNB1 | 0.0020 | linker gene; AJ component |
| Protein interactions – continued | | |
| DIAPH1 - RHOC | 0.0020 | DEG |
| CTNNB1 - CTNNA2 | 0.0020 | AJ components |
| JUP - CTNNA2 | 0.0020 | linker gene; AJ component |
| PVR - NECTIN3 | 0.0019 | linker gene; AJ component |
| CDK2 - LIG1 | 0.0017 | DEG |
| CDK2 - CCNA2 | 0.0016 | DEG |
| NECTIN4 - NECTIN1 | 0.0016 | AJ components |
| CTNNB1 - CDH1 | 0.0014 | AJ components |
| PKMYT1 - CDK1 | 0.0014 | DEG |
| JUP - CTNNA1 | 0.0014 | linker gene; AJ component |
| CDK2 - CKS1B | 0.0014 | DEG |
| CDK1 - CCNA2 | 0.0014 | AJ components |
| NECTIN3 - NECTIN1 | 0.0013 | AJ components |
| CTNNB1 - CTNNA1 | 0.0013 | AJ components |
| NECTIN4 - NECTIN2 | 0.0010 | AJ components |
| PTRM - CTNND1 | 9.89E-04 | DEG; linker genes |
| CDK2 - RBL1 | 9.26E-04 | DEG |
| PKMYT1 - CCNB2 | 7.25E-04 | DEG |
| CCNB2 - CKS1B | 6.15E-04 | DEG |
| CTNNB1 - CTNNA3 | 5.53E-04 | AJ components |
| CCNB1 - CCNB2 | 4.81E-04 | DEG |
| CTNNA1 - CTNNA3 | 4.77E-04 | AJ components |
| AFDN - NECTIN4 | 4.19E-04 | AJ components |
| CCNB1 - CDK2 | 3.86E-04 | DEG |
| JUP - CTNNA3 | 3.76E-04 | linker gene; AJ component |
| ELF3 - EPS15 | 3.39E-04 | DEG; linker genes |
| CCNB1 - CKS1B | 3.22E-04 | DEG |
| PKMYT1 - CCNB1 | 3.18E-04 | DEG |
| Protein interactions – continued | | |
| AFDN - NECTIN3 | 3.10E-04 | AJ components |
| AFDN - NECTIN1 | 2.55E-04 | AJ components |
| CKS1B - CCNA2 | 2.50E-04 | DEG |
| TCF7L2 - JUP | 2.46E-04 | linker gene; AJ component |
| PKMYT1 - CCNA2 | 2.44E-04 | DEG |
| AFDN - NECTIN2 | 2.25E-04 | AJ components |
| ECT2 - GPSM2 | 2.13E-04 | DEG |
| CCNA2 - RBL1 | 1.57E-04 | DEG |
| CDH1 - ARHGAP21 | 1.49E-04 | DEG; AJ component |
| MYH10 - LGALS3 | 1.28E-04 | DEG |
| CTNNA1 - CTNND1 | 1.21E-04 | AJ components |
| RBL1 - E2F2 | 1.15E-04 | DEG |
| CTNNB1 - RACGEF2 | 1.13E-04 | DEG; AJ component |
| CTNNA1 - AFDN | 1.11E-04 | AJ components |
| CCNB1 - CCNA2 | 1.06E-04 | DEG |
| ELF3 - EGFR | 1.01E-04 | DEG; linker genes |
| CDK1 - ECT2 | 9.21E-05 | DEG |
| JUP - CTNND1 | 9.18E-05 | DEG; linker genes |
| CCNA2 - LMNB1 | 8.37E-05 | DEG |
| CDH1 - ECT2 | 7.60E-05 | DEG; AJ component |
| JUP - CTNNB1 | 7.27E-05 | linker gene; AJ component |
| PTRM - CTNNB1 | 6.11E-05 | linker gene; AJ component |
| CDK1 - RACGAP1 | 6.03E-05 | AJ components |
| JUP - CDH1 | 5.93E-05 | linker gene; AJ component |
| RACGAP1 - STON2 | 5.80E-05 | DEG |
| JUP - ERBB2 | 5.67E-05 | DEG; linker genes |
| CCNB1 - LMNB1 | 5.64E-05 | DEG; linker genes |
| SP1 - MYH10 | 5.63E-05 | DEG; linker genes |
| Protein interactions – continued | | |
| PVR - CDH1 | 5.37E-05 | linker gene; AJ component |
| CDK1 - LMNB1 | 4.39E-05 | AJ components |
| MYH10 - ARHGAP21 | 4.35E-05 | DEG |
| MYH10 - STON2 | 4.26E-05 | DEG |
| EPS15 - MYH10 | 4.25E-05 | DEG; linker genes |
| SP1 - ARHGAP21 | 4.17E-05 | DEG; linker genes |
| SP1 - RBL1 | 3.77E-05 | DEG; linker genes |
| CDK2 - LMNB1 | 3.68E-05 | DEG |
| RACGAP1 - RACGEF2 | 3.30E-05 | DEG |
| LMNA - CDH1 | 3.04E-05 | linker gene; AJ component |
| LMNA - CDK1 | 2.64E-05 | DEG; linker genes |
| EPS15 - EGFR | 2.44E-05 | linker genes |
| CTNNB1 - LMNB1 | 2.24E-05 | DEG; AJ component |
| CCNB1 - CDH1 | 2.21E-05 | DEG; AJ component |
| EPS15 - ERBB2 | 2.14E-05 | DEG; linker genes |
| CTNNB1 - CTNND1 | 2.13E-05 | AJ components |
| ERBB2 - CTNNB1 | 1.88E-05 | DEG; AJ component |
| EGFR - ERBB2 | 1.87E-05 | DEG; linker genes |
| SP1 - CCNA2 | 1.52E-05 | DEG; linker genes |
| CTNNB1 - CCNA2 | 1.34E-05 | DEG; AJ component |
| CDK2 - RHOC | 1.33E-05 | DEG |
| LMNA - CTNNB1 | 1.25E-05 | linker gene; AJ component |
| JUP - MYH10 | 1.25E-05 | DEG; linker genes |
| SP1 - CCNB1 | 1.02E-05 | DEG; linker genes |
| EGFR - LGALS3 | 9.97E-06 | DEG; linker genes |
| LMNA - RACGEF2 | 8.47E-06 | DEG; linker genes |
| EGFR - CTNNB1 | 7.32E-06 | linker gene; AJ component |
| SP1 - CDK1 | 7.13E-06 | DEG; linker genes |
| Protein interactions – continued | | |
| SP1 - CDK2 | 6.68E-06 | DEG; linker genes |
| CDK2 - CTNNB1 | 5.90E-06 | DEG; AJ component |
| CDK2 - MYH10 | 4.24E-06 | DEG |
| SP1 - CTNNB1 | 4.08E-06 | linker gene; AJ component |
| EGFR - CTNNA1 | 2.89E-06 | linker gene; AJ component |
| EGFR - EXOC5 | 2.63E-06 | DEG; linker genes |
| CDK2 - JUP | 2.32E-06 | DEG; linker genes |
| LMNA - CDK2 | 1.27E-06 | DEG; linker genes |
| CDK2 - CDK1 | 1.21E-06 | DEG |
