## Supplementary figures and images for "CG7379/ING1 suppress cancer cell invasion by maintaining cell-cell junction integrity"

### Supplemental Figures

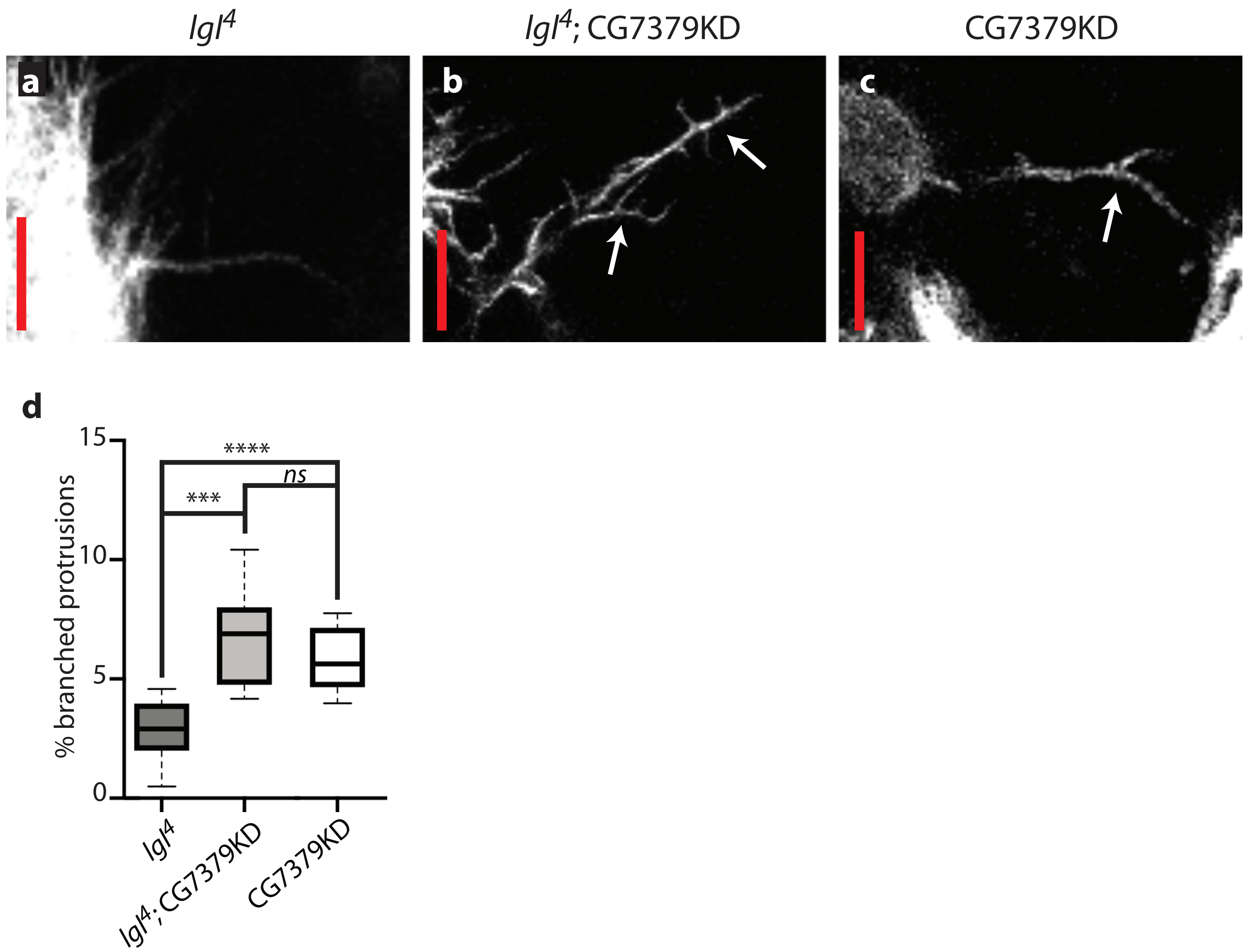

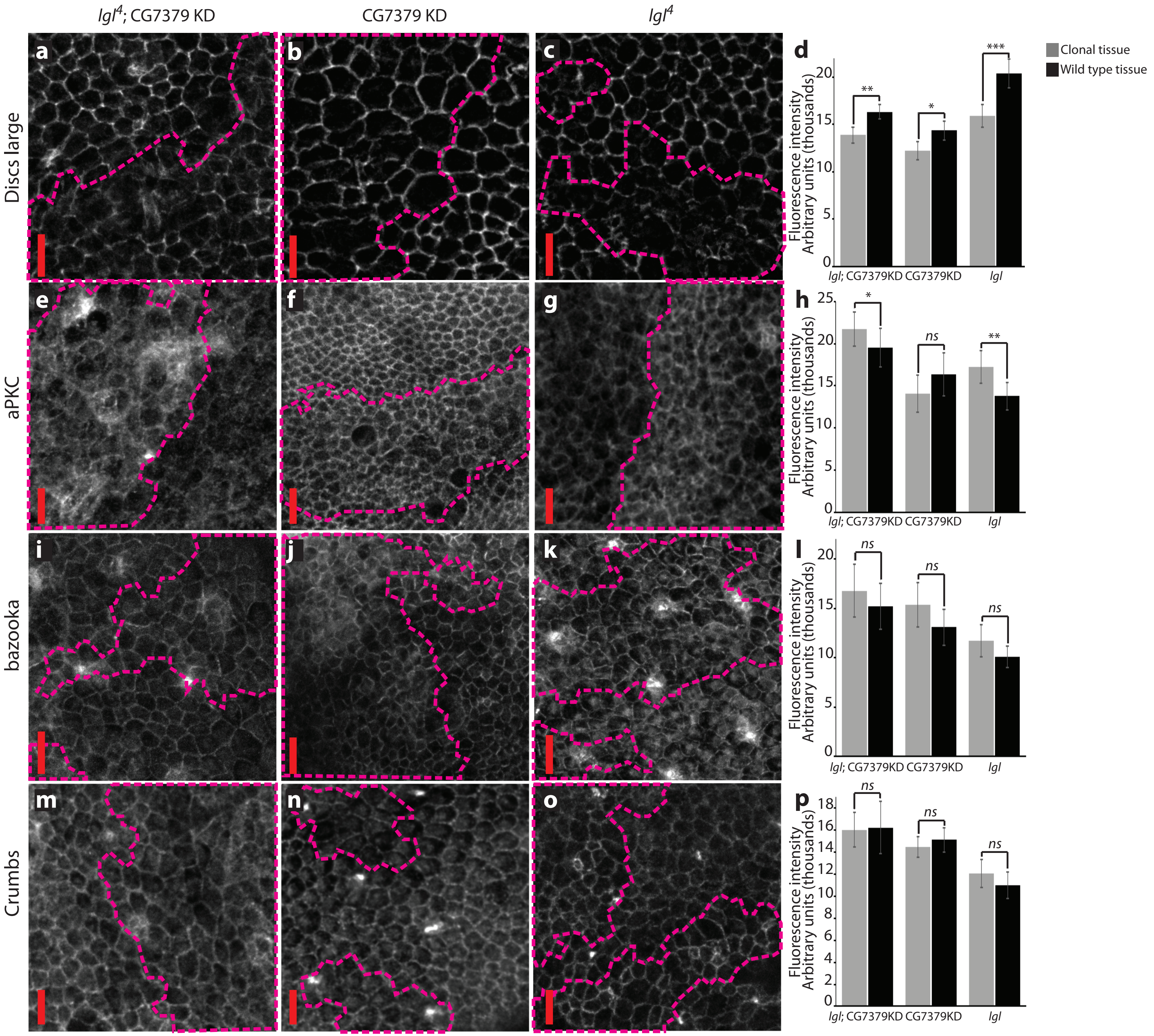

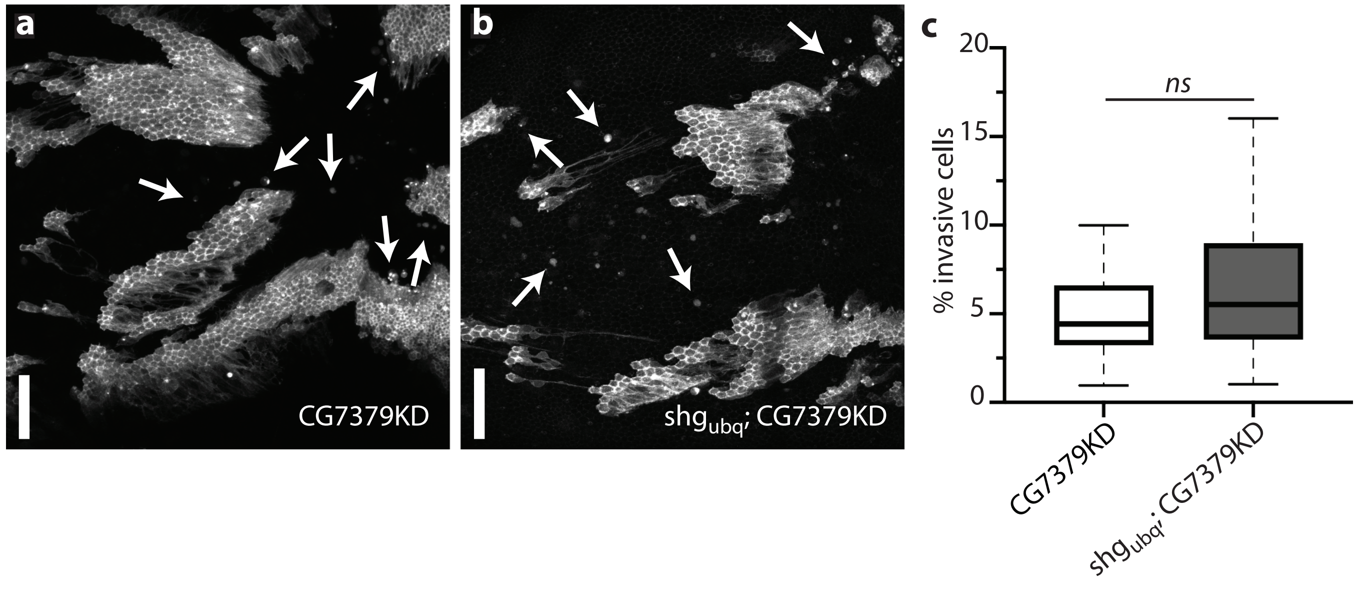

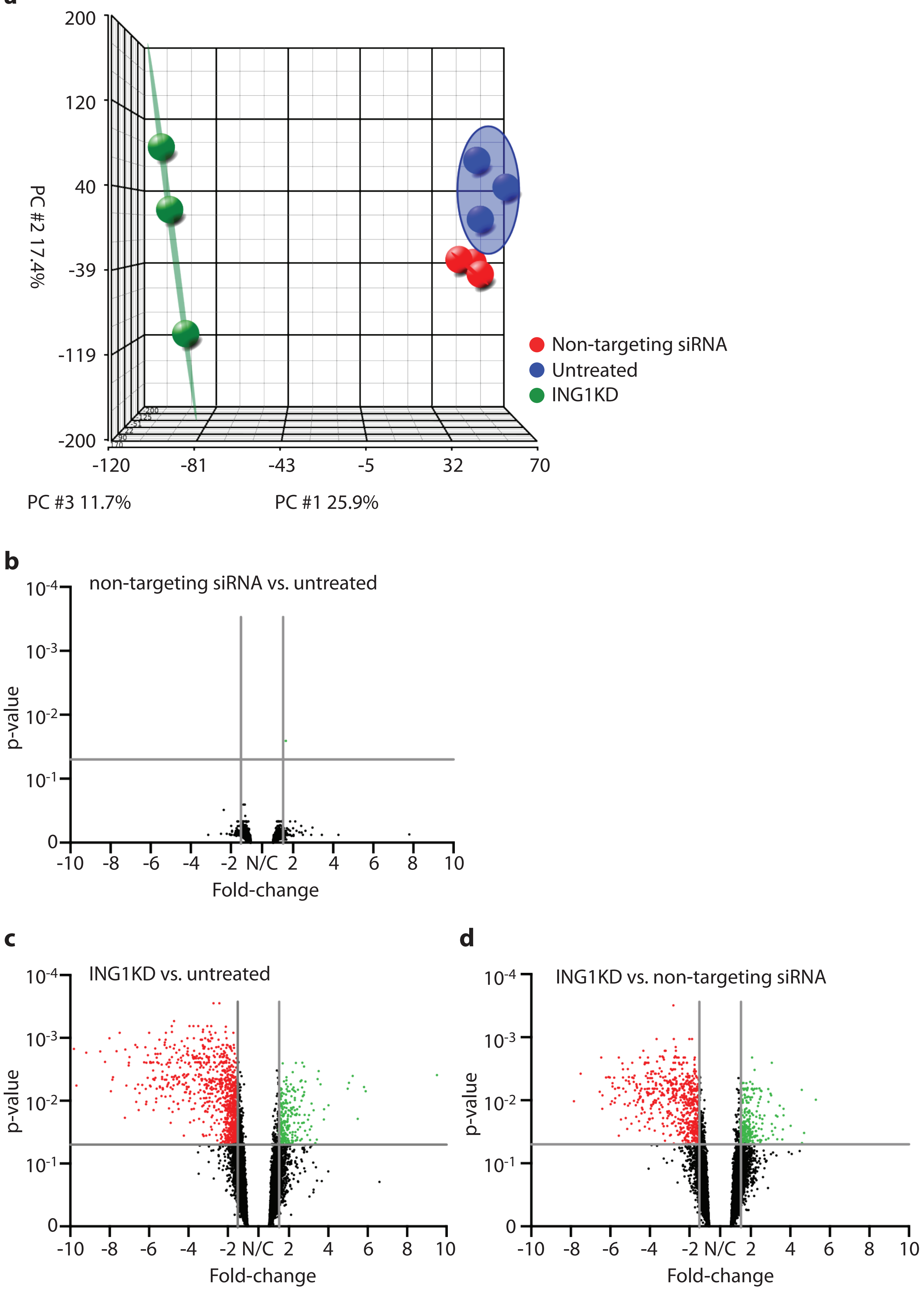
